# Gut bacteria generate prodrugs in situ increasing systemic drug exposure

**DOI:** 10.64898/2025.12.09.692405

**Authors:** Ting-Hao Kuo, Anoop Singh, Amber Brauer-Nikonow, Mahnoor Zulfiqar, Resul Gökberk Elgin, Li-Yao Chen, Matthias Gross, George-Eugen Maftei, Michael Zimmermann

## Abstract

The human gut microbiome has been increasingly recognized as a metabolic compartment that influences health and therapeutic responses. While recent studies have cataloged diverse sets of drug biotransformation reactions as a result of microbial metabolism^1–3^, the pharmacological relevance of the resulting drug metabolites remains mostly uncharacterized. We investigated whether certain microbial drug metabolites can act as functional prodrugs. By systematically mining 871 putative drug metabolites produced by gut bacteria^1^, we identified microbial drug metabolites with key physicochemical properties of prodrugs. Using bacterial culturing, a genetic gain-of-function screen coupled to mass spectrometry and comparative genomics, we uncover that *Bacteroidota* bacteria encode conserved methyltransferases that can generate prodrugs through carboxyl methylation. As a case in point, we demonstrate that bezafibrate gets converted to a prodrug by a distinct bacterial enzyme, increasing epithelial drug permeability *in vitro* and systemic drug exposure *in vivo* in a gnotobiotic mouse model. Additionally, we tested 170 structurally and clinically diverse drugs, and demonstrate that microbial drug–prodrug conversion is a common result of gut bacterial drug biotransformation. Altogether, our findings suggest a novel mechanism of how the gut microbiota influences an individual’s pharmacokinetics, and may generally cause interpersonal differences in microbiota–host metabolic interactions.

## Main

Prodrug design is a cornerstone of modern pharmacology. By conjugating bioreversible chemical moieties, otherwise sub-optimal oral drugs gain improved oral bioavailability and systemic exposure, leading to more favorable pharmacokinetic profiles^4,5^. Such prodrugs are pharmacologically inactive and are typically converted by human enzymes, most prominently hydrolases^6^ and cytochrome P450 enzymes^7^, to regenerate the active parent drug following intestinal absorption.

In the human gut, trillions of microorganisms collectively form a microbial ecosystem that drives complex chemical transformations of drug molecules and other compounds^1,3,8,9^. Some of these microbial metabolic activities have been suggested to contribute to inter-individual differences in pharmacokinetics and drug responses^10–12^, for example through drug toxification, (in)activation, and by modulating enterohepatic drug circulation^13–15^. Recent studies employing large-scale microbiota–drug metabolic biotransformation assays have revealed previously unrecognized microbial activity on drug metabolism and resulting metabolites^1–3^, for most of which the functional consequences however remained underexplored.

We hypothesized that certain gut microbial drug metabolites can function as prodrugs and thereby enhance systemic exposure of the parent drug. We indeed identified microbial drug metabolites bearing key physicochemical properties of prodrugs (namely, increased molecular mass and hydrophobicity) among a set of 871 putative bacterial drug biotransformation products. Experimental validation of drug metabolite structures, identification of biotransformation enzymes, and *in vitro* and *in vivo* demonstration of prodrug properties suggests microbial prodrug formation as an underappreciated determinant of drug disposition and introduce a new perspective on microbiome-informed pharmacology.

### Mining microbial drug metabolites for putative prodrugs

To screen microbial drug metabolites with prodrug potential, we leveraged a metabolomics dataset of gut microbial metabolism of therapeutic drugs previously reported (ChEMBL ID: CHEMBL5303573; MetaboLights accession: MTBLS896)^1^. This dataset comprises 19,783 metabolic microbe-drug interactions, generated through incubation of 271 oral drugs with 73 representative human gut bacterial strains under anaerobic conditions, and analyzed with reversed-phase liquid chromatography (RPLC) coupled to high-resolution mass spectrometry (LC–MS) (Fig. 1a). Untargeted metabolomics identified 871 putative metabolites from 176 biotransformed drugs, spanning a broad range of mass differences (Δmass) and chromatographic retention time shifts (ΔRT) relative to their respective parent drugs (Fig. 1b; Supplementary Table 1).

**Fig. 1.**
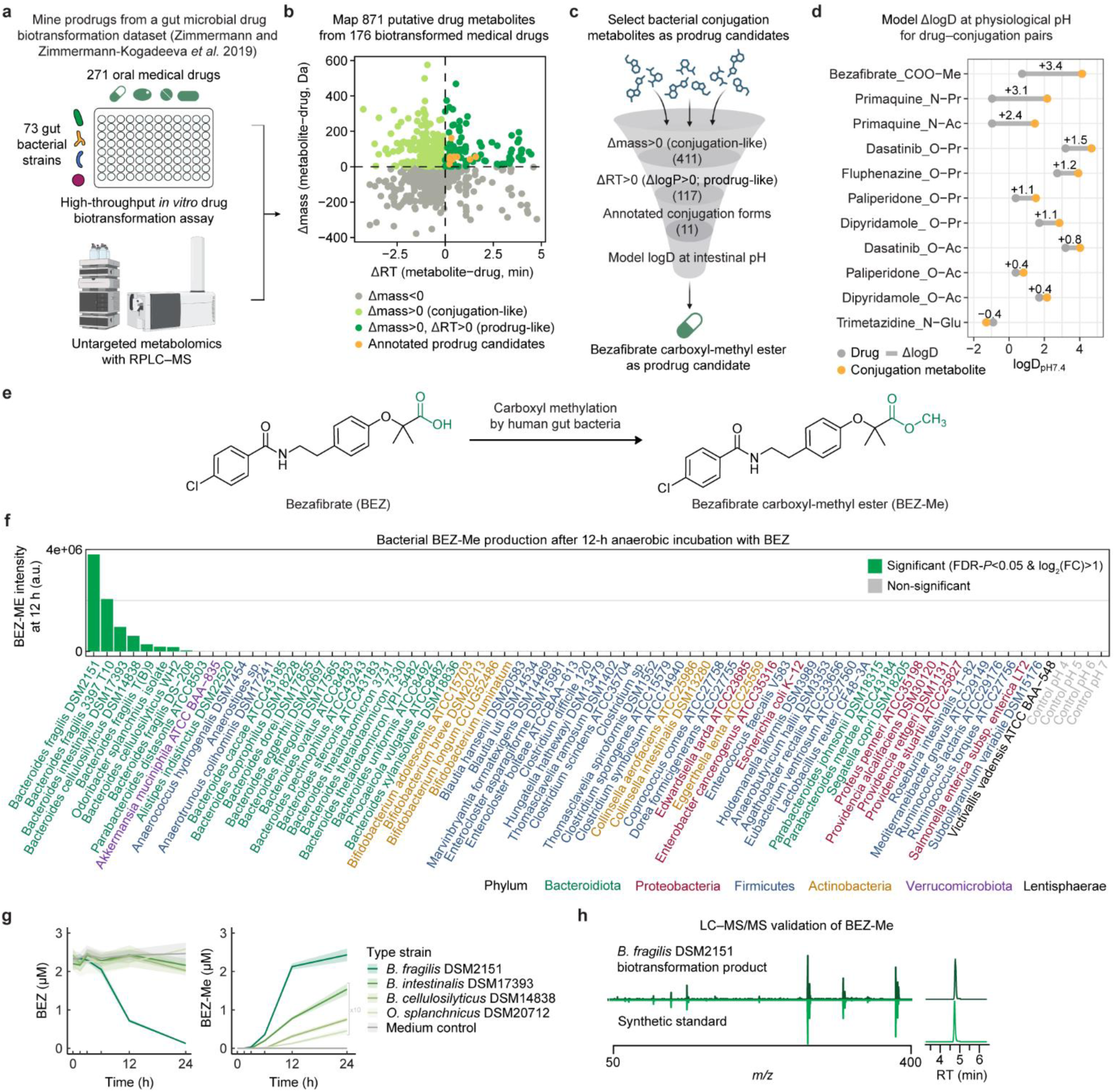
Mine bacterial drug metabolites for prodrug candidates. **a**, Overview of the previously reported study on gut microbial drug metabolism for mining functional bacterial drug metabolites^1^. **b**, Mapping of 871 putative bacterial drug metabolites by differences in RT and molecular mass compared to their respective parent drugs (Supplementary Table 1). Grey, 450 metabolites with loss in mass; light green, 411 conjugation-like metabolites with Δmass > 0 and ΔRT < 0; dark green, 117 prodrug-like metabolites with Δmass > 0 and ΔRT > 0, among which 11 with resolved conjugation forms are identified (orange; Supplementary Table 2). **c**, Scheme for selecting bacterial drug metabolites as prodrug candidates. **d**, Modeling logD at pH 7.4 between each prodrug candidate and its parent drug (Supplementary Table 2). Conjugation form of each metabolite is indicated (‘drug name_conjugation form’, y-axis). The number denoted for each pair indicates ΔlogD = logD_prodrug_ – logD_drug_. **e**, Reaction scheme: BEZ conversion to BEZ-Me. **f**, Reported BEZ-methylating activity in gut bacteria^1^. Bars represent the mean LC–MS peak area of BEZ-Me at 12 h of bacterial incubation with BEZ (2 µM), calculated from 4 independent cultures (Supplementary Table 3). Statistical significance for comparisons between 12 h and 0 h was determined by FDR-adjusted *P* < 0.05 and log_2_(fold change) > 2. **g**, *In vitro* BEZ methylation assay of selected type strains from the active species in panel (f) (2 µM of BEZ as substrate; Supplementary Table 4). Line and shaded area depict mean and s.d., respectively, calculated from 4 independent assay replicates. **h**, Structural confirmation of the bacterial biotransformation product of BEZ as BEZ-Me by LC–MS/MS and comparison to chemical standard. LC–MS and drug funnel symbols were created with BioRender.com.

To identify microbial drug metabolites with prodrug potential, we sequentially filtered the metabolites based on their key physiological properties resembling those of prodrugs (Fig. 1c). We first selected 411 of the 871 metabolites with a positive Δmass value, reflecting a fundamental structural characteristic of prodrugs bearing a conjugation moiety (Fig. 1b,c). In line with the primary goal in prodrug design to boost molecular hydrophobicity and to improve membrane permeability^16^, we prioritized metabolites with positive ΔRT values, an empirical proxy for increased hydrophobicity (Supplementary Fig. 1a). This yielded 117 microbial drug metabolites with prodrug-like physiochemical properties (Fig. 1b,c) for further investigations.

To gain structural insights into these putative prodrug candidates, we subsequently mapped Δmass values to known drug conjugation reactions^17^ allowing to propose the chemical structure of 11 drug metabolites derived from 7 distinct parent drugs (orange dots in Fig. 1b; Supplementary Table 2). These putative drug metabolites resulted from different conjugation reactions including methylation (Me, CH_2_, +14.012 Da), acetylation (Ac, C_2_H_2_O, +42.012 Da), propionylation (Pr, C_3_H_4_O, +56.026 Da), and glycosylation (Glu, C_6_H_10_O_5_, +162.053 Da). The presence of a conjugatable functional group in each parent drug, such as a carboxyl group (-COOH), a primary or secondary amine (-NH_2_ or -NH-), or a hydroxyl group (-OH), further supports the proposed conjugation of the respective parent drugs (Supplementary Fig. 1b,c). To assess the extent to which the conjugation of a given drug could enhance intestinal drug permeability, we modeled pH-dependent lipophilicity (logD) at physiological pH 7.4^18^ for each of the 11 drug–conjugate pairs (Fig. 1d; Supplementary Table 2). Among all drug–metabolite pairs examined, the methylation of bezafibrate (BEZ), a lipid-lowering drug to treat hyperlipidaemia^19^, into bezafibrate carboxyl-methyl ester (BEZ-Me) resulted in the largest increase in hydrophobicity (ΔlogD_pH7.4_ = 3.4) (Fig. 1e). The resulting high lipophilicity (logD_pH7.4_ = 4.1) of BEZ-Me suggests that carboxyl methylation substantially enhances membrane permeability, as well as potential oral bioavailability, compared to BEZ. These prodrug-like physicochemical properties of BEZ-Me are in agreement with industrially developed fibrate prodrugs (*e.g.*, fenofibrate^20^) as well as several clinically successful carboxyl-methyl ester prodrugs^21–23^. Therefore, we further investigated BEZ and sought to characterize its microbial biotransformation mechanisms and to evaluate its capability of enhancing intestinal absorption and systemic drug distribution.

Among the 73 bacterial strains previously assayed for drug biotransformation, we identified 8 strains belonging to 4 different species within the *Bacteroidota* phylum *(Bacteroides fragilis, Bacteroides cellulosilyticus, Bacteroides intestinalis*, and *Odoribacter splanchnicus*) that demonstrated marked BEZ methylation activity (Fig. 1f; Supplementary Table 3). To validate these previous findings, we anaerobically incubated BEZ with the type strains of each of the four active species, all of which successfully produced BEZ-Me (Fig. 1g; Supplementary Table 4). We also confirmed the chemical structure of bacterially produced BEZ-Me by liquid chromatography tandem mass spectrometry (LC–MS/MS) in comparison with the standard compound that we chemically synthesized (Fig. 1h; Supplementary Data).

### Drug methyltransferases producing prodrugs are widely distributed among gut bacteria

To identify microbial enzymes that convert BEZ to BEZ-Me, we employed a genetic gain-of-function (GoF) approach^1^ using *B. fragilis* DSM2151 (the strain with the strongest BEZ methylating activity, Fig. 1e) as DNA source (Fig. 2a). In brief, we generated heterologous *Escherichia coli* clones with inserted genomic fragments of *B. fragilis* DSM2151 and subsequently tested their gained capacity to perform BEZ methylation. The resulting library contained 36,480 picked and arrayed *E. coli* clones, each heterologously expressing a ∼2–4 kb genomic fragment (average size ∼2.4 kb), representing an 18-fold coverage of the *B. fragilis* genome (Supplementary Fig. 2a,b). We incubated the library with BEZ (1.5 µM) anaerobically, applied high-throughput LC–MS measurements to detect the production of BEZ-Me, and identified one active clone with a genomic fragment encoding two genes, *bf2169* and *bf2170* (Fig. 2b; Supplementary Fig. 2c,d). Heterologous expression of either of the two open reading frames in *E. coli* and testing for BEZ–to–BEZ-Me conversion pinpointed *bf2170*, encoding a putative S-adenosyl-methionine-dependent methyltransferase, as gene underlying the observed methylation of BEZ (Supplementary Fig. 2e; Supplementary Table 5). To further corroborate these findings, we made a *bf2170*-deletion mutant (Δ*bf2170*) and complementations at different gene expression levels in *B. fragilis* DSM2151, and demonstrated that *bf2170* is necessary and sufficient for BEZ methylation (Fig. 2c; Supplementary Table 6), whereas deletion of *bf2170* did not affect bacterial growth (Supplementary Fig. 2f).

**Fig. 2.**
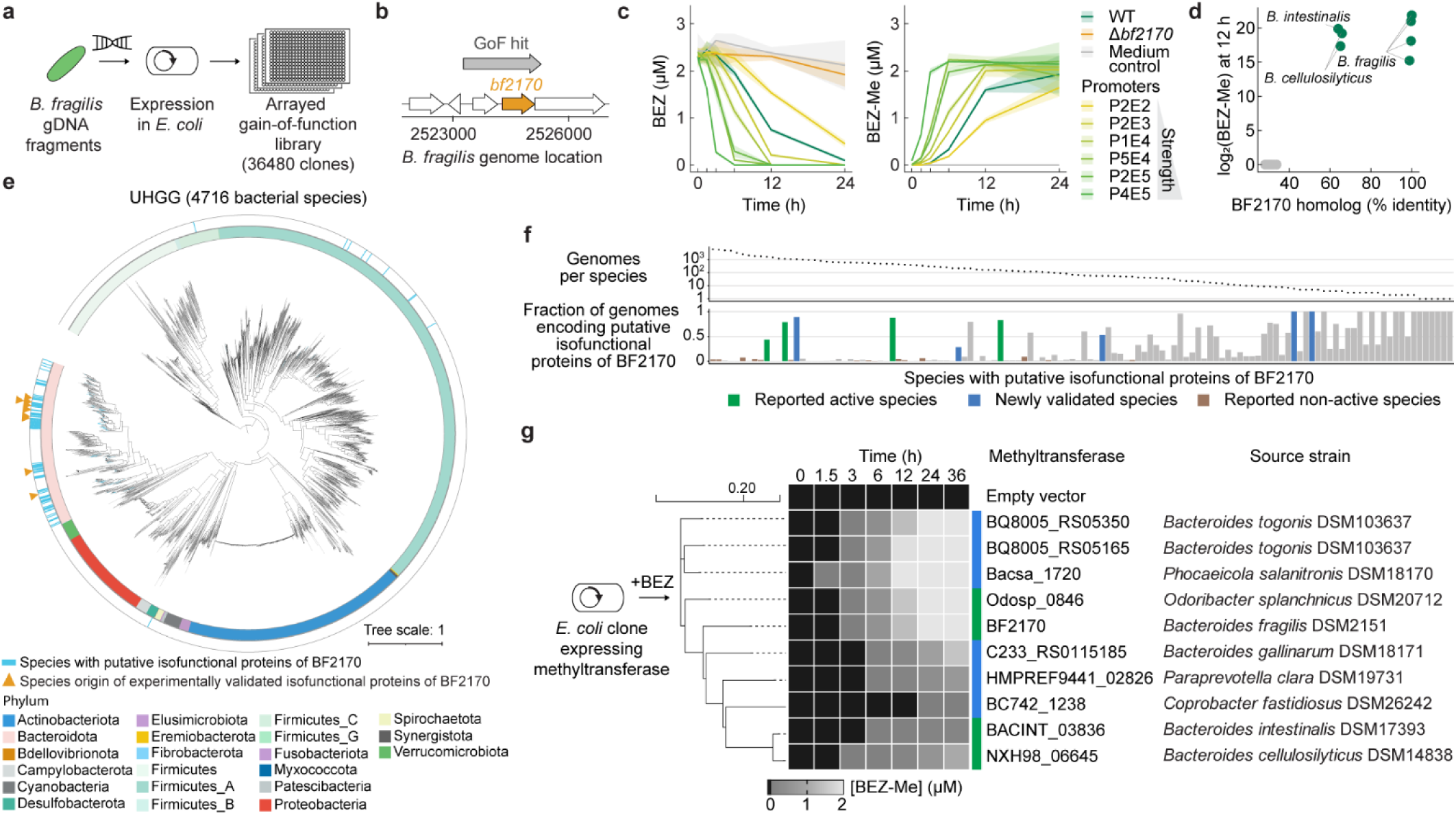
Identify and characterize *Bacteroidota* methyltransferases generating the bezafibrate prodrug. **a**, The GoF approach for identifying gut bacterial enzymes methylating BEZ. Genomic DNA from *B. fragilis* DSM2151 was sheared, heterologously cloned into *E. coli*, arrayed, and tested for BEZ-methylating activity. **b**, Mapping a methylation-active GoF insert sequence to the *B. fragilis* genome showed a gene locus encoding BF2170, a putative methyltransferase. **c**, BEZ-methylating activity of *B. fragilis* WT, Δ*bf2170* mutant, and *bf2170*-complemented strains at different expression levels (P4E5 > P2E5 > P5E4 > P1E4 > P2E3 > P2E2)^1,36^, assessed by *in vitro* anaerobic assays (2 µM of BEZ as substrate; Supplementary Table 6). Line and shaded area depict mean and s.d., respectively, calculated from 4 independent assay replicates. **d**, Correlation between BEZ-methylating activity and the presence of BF2170 homolog among the 69 strains with publicly available genomes of the 73 strains shown in Fig. 1f (Supplementary Table 7). **e**, Phylogenetic tree of 4,716 representative bacterial species from the UHGG collection. Blue bar (outer ring), putative isofunctional proteins of BF2170 distributed among 124 bacterial species (Supplementary Tables 8, 9, and 10); orange triangle, species origin of BF2170 and 9 additional methyltransferases experimentally validated in this study (Supplementary Table 11). **f**, Fraction of genomes encoding putative isofunctional proteins of BF2170 in each hit species (Supplementary Table 10; see Supplementary Fig. 3b for detailed legends). **g**, Functional validation of 9 selected *Bacteroidota* methyltransferases (orange triangles in panel (e); green and blue bars in panel (f)) by heterologous expression in *E. coli* and *in vitro* anaerobic assays (2 µM of BEZ as substrate). Clones expressing BF2170 or an empty vector were included as controls. Methyltransferases are clustered by multiple sequence alignment. Tile color depicts mean BEZ-Me concentration, quantified from 4 independent assay replicates (Supplementary Table 12).

To investigate whether the identified *B. fragilis* gene, *bf2170*, is representative of the BEZ methylation activity in the gut microbiota, we first searched for BF2170 homologs across the 73 bacterial strains previously tested for BEZ biotransformation^1^ (Fig. 1f). Indeed, the genomic presence of isofunctional proteins of BF2170 (protein sequence identity > 63.9%) discriminated between strains producing BEZ-Me and all the other strains (Fig. 2d; Supplementary Table 7). To gain insights into the taxonomic distribution of isofunctional proteins of BF2170 across the human gut microbiome, we then queried the Unified Human Gastrointestinal Genome (UHGG) collection^24^, encoding >170 million protein sequences from 289,232 prokaryotic genomes. This analysis identified 4,937 putative isofunctional proteins of BF2170 from 4,794 bacterial genomes (protein sequence identity > 61%; Supplementary Table 8 and 9) distributing across 124 (2.6%) out of the 4,716 representative bacterial species in the UHGG (Fig. 2e; Supplementary Table 10). Most of these hits (92%; 114/124 species) belong to the *Bacteroidiota* phylum (Supplementary Fig. 3a), with substantial variation in strain-level prevalence (Fig. 2f; Supplementary Fig. 3b). The majority of the genomes contain only a single hit for putative isofunctional proteins of BF2170 (97.1 %; 4,656/4,794), whereas a small fraction of the genomes carries two (2.8%; 133/4,794) or three (0.1%; 5/4,794) (Supplementary Fig. 3c; Supplementary Table 8). To functionally validate these putative genes, we heterologously expressed 9 additional methyltransferases from 8 distinct species of the *Bacteroidiota* phylum in *E. coli* (Fig. 2f; Supplementary Table 11), assessed their methylation activity, and found that all tested methyltransferases are capable of methylating BEZ, however with varying biotransformation efficiency across the enzymes (Fig. 2g; Supplementary Table 12). Since these methyltransferases exhibited high similarity in predicted overall protein structures (RMSD = 0.398 Å; Supplementary Fig. 3d; Supplementary Table 13), subtle residue-level variations within the catalytic pocket might likely account for the observed differences in methylation activity. Overall, these findings establish that certain gut bacteria of the *Bacteroidota* phylum carry genes encoding a sequence–conserved methyltransferase that can generate the methyl prodrug of BEZ.

### Bacterial bezafibrate methylation facilitates intestinal absorption

Having identified BEZ-Me as a putative prodrug produced by specific bacterial members of the gut microbiota, we sought to compare its intestinal permeability to its parent drug, BEZ. To this aim, we employed a Caco-2 transwell assay, a well-established *in vitro* model to quantify trans-epithelial permeability of drugs^25^. We applied BEZ or an equimolar amount of BEZ-Me to the apical side, and quantified their concentration kinetics on both the apical and basolateral sides using LC–MS (Fig. 3a–c; Supplementary Table 14). As expected, BEZ alone showed poor permeability with only ∼1% of the applied dose appearing on the basolateral side (Fig. 3a) and a low apical-to-basolateral apparent permeability coefficient (P_app_ = (1.3 ± 0.13) × 10^-9^ cm/s; Fig. 3c). In contrast, BEZ-Me enters the cells, and is de-methylated producing a ∼5-fold increase in basolateral BEZ levels (Fig. 3b) and significantly enhanced BEZ permeability (P_app_ = (2.8 ± 0.65) × 10^-8^ cm/s; Student’s *t*-test, *P* = 5.5 × 10^-7^; Fig. 3c; Supplementary Table 15). The observed trans-epithelial hydrolysis of BEZ-Me is consistent with the canonical bioactivation mechanism of ester prodrugs, wherein ubiquitous cellular hydrolases cleave ester moieties (de-methylation in this case) to release the active carboxyl drug^26^. Reversing the assay orientation by dosing from the basolateral side altered the transport kinetics, suggesting the involvement of directional transport mechanisms^27^ (Supplementary Fig. 4a–c; Supplementary Table 14). The transport profile was reproducible at a 20-fold lower dose (Supplementary Fig. 4e–i; Supplementary Table 16). Altogether, these results propose a hydrolysis-coupled trans-epithelial bioactivation of BEZ-Me and suggest that intestinal BEZ-Me could indeed enhance BEZ absorption in a prodrug-like manner.

**Fig. 3.**
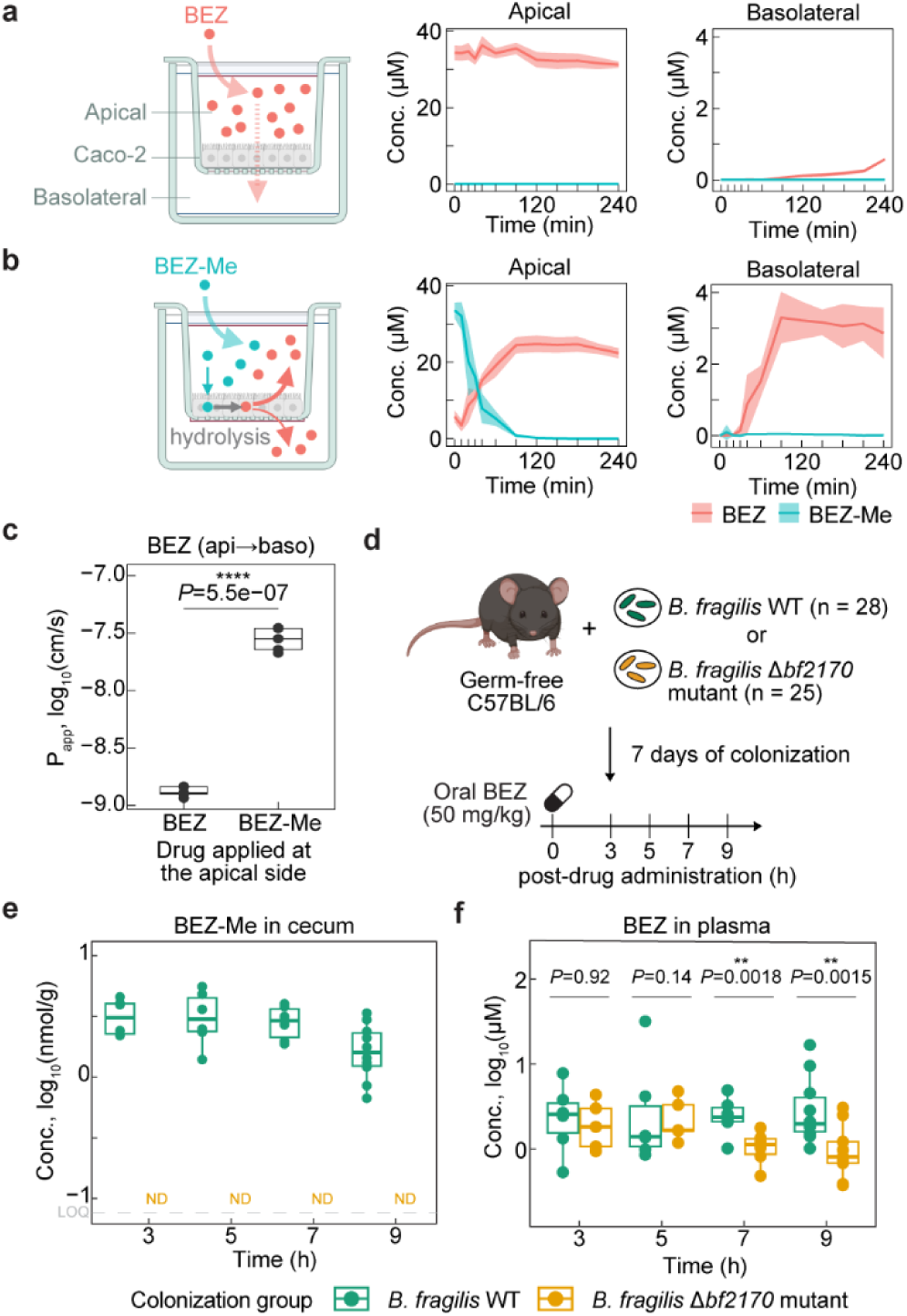
Test prodrug properties of bezafibrate methyl ester *in vitro* and in a gnotobiotic mouse model. **a** and **b**, Caco-2 transwell assay for assessing trans-epithelial permeability of BEZ. BEZ (40 µM) (a) or BEZ-Me (40 µM) (b) was applied to the apical side, and the concentration kinetics of BEZ and BEZ-Me in both apical and basolateral sides was quantified over 4 h. Line and shaded area depict mean and s.d., respectively, calculated from 5 independent assay replicates (Supplementary Table 14). **c**, P_app_ of BEZ in the apical-to-basolateral direction obtained from (a) and (b). Pairwise comparison was performed using a two-sided Student’s *t*-test with unequal variance (**** *P* < 0.0001; Supplementary Table 15). **d**, Gnotobiotic mouse model for studying the pharmacokinetics of BEZ. Germ-free C57BL/6 mice mono-colonized with either *B. fragilis* WT (*n* = 28) or Δ*bf2170* mutant (*n* = 25) were orally administered a single dose of oral BEZ (50 mg/kg of body weight) and samples were collected at 3, 5, 7, or 9 h post drug administration (see Supplementary Table 17 and ‘*Statistics and reproducibility*’ in Methods for group and batch details). **e**, Cecum concentration of BEZ-Me (Supplementary Table 18). LOQ: limit of quantification; ND: not detected. **f**, Plasma concentration of BEZ. Pairwise comparisons between colonization groups at each time point were based on estimated marginal means from a two-way linear model and tested with two-sided Tukey’s test (** *P* < 0.01; Supplementary Table 19). For all box plots, boxes indicate the interquartile range (25^th^–75^th^ percentiles) with the centre line denoting the median; whiskers extend to data points within 1.5× IQR from the hinges. Transwell and mouse symbols were created with BioRender.com.

### Bacterially produced bezafibrate prodrug enhances systemic drug exposure in a gnotobiotic model

To investigate whether the bacterial conversion of BEZ to BEZ-Me affects pharmacokinetics, we took advantage of the generated genetic Δ*bf2170* strain of *B. fragilis* DSM2151 to establish a gnotobiotic mouse model. We mono-colonized germ-free (GF) mice with either the wild-type strain (GF^WT^) or the Δ*bf2170* mutant (GF^Δ*bf2170*^) (n = 28 and 25, respectively), followed by oral administration of a single dose of BEZ (50 mg/kg) with a treatment duration of 3, 5, 7, or 9 h (Fig. 3c; Supplementary Table 17). As expected from the *in vitro* data, we detected BEZ-Me solely in the large intestine of GF^WT^ mice, but not in GF^Δ*bf2170*^ mice (Fig. 3d; Supplementary Table 18), demonstrating that BEZ reaches the large intestine and that BF2170 is expressed and active in the intestinal environment. Next, we asked whether bacterial methylation of BEZ in the intestine increases systemic exposure of BEZ. Indeed, we found that the plasma level of BEZ in GF^WT^ mice is significantly higher than in GF^Δ*bf2170*^ mice at 7 and 9 h post oral administration (Tukey’s post hoc test, *P* = 0.018 and 0.0015 at 7 and 9 h, respectively) (Fig. 3e; Supplementary Table 19). The observed difference only at later time points is explained by the difference in timing between the fast small intestinal absorption of BEZ compared to the delayed contribution of bacterially produced BEZ-Me to the pharmacokinetics^11^. Altogether, these data functionally validate the predicted prodrug property of BEZ-Me and the hypothesized capacity of gut bacteria to alter drug pharmacokinetics by biotransforming drugs to prodrugs.

### Gut bacteria methylate chemically diverse carboxyl drugs

To test whether the identified gut bacterial production of methyl ester prodrugs also applies to other drugs, we screened 170 carboxyl drugs using an *E. coli* heterologous expression clone producing BF2170 as a representative enzyme under anaerobic conditions (Fig. 4a). These drugs span a broad chemical space when mapped to small-molecule drug entities in DrugBank^28^ (Fig. 4b) and encompass diverse chemical structures and Anatomical Therapeutic Chemical (ATC) therapeutic classes (Fig. 4c; Supplementary Table 20). Besides BEZ, we identified 14 additional drugs that yielded methylation products (orange dots in Fig. 4b and highlighted in Fig. 4c), 13 of which were structurally confirmed by LC–MS/MS comparison to standards that we chemically synthesized and purchased (Fig. 4d; see Supplementary Data for LC–MS/MS spectra). BF2170 preferentially methylates fibrate drugs used to manage dyslipidemia, including BEZ, efaproxiral (a BEZ analogue), and fenofibric acid, while also acting on profen-class nonsteroidal anti-inflammatory drugs (ketoprofen, zaltoprofen, pranoprofen, indobufen, tiaprofenic acid, bromfenac, and ketorolac) and on a few other drug compounds (cardarine, Ki16425, timapiprant, orantinib, and nateglinide), reflecting broad substrate promiscuity. These results suggest that the *Bacteroidota*-encoded methyltransferases can convert chemically diverse carboxyl drugs into their prodrugs, suggesting microbial prodrug production as a general route through which the gut microbiota can affect systemic drug exposure.

**Fig. 4.**
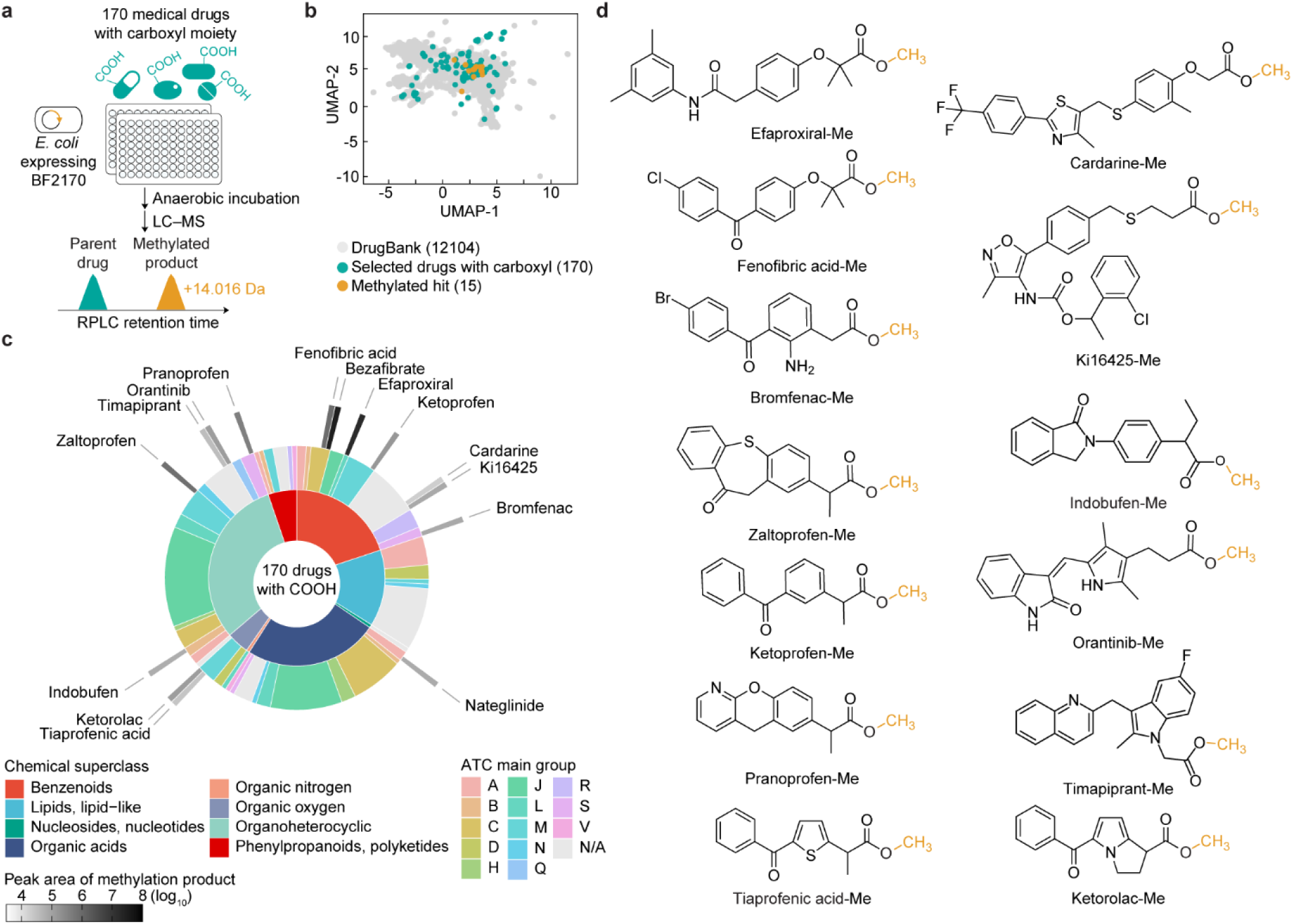
Screen additional drug–prodrug conversion by gut bacterial methyltransferase. **a**, High-throughput screening of drug–prodrug conversion by the bacterial methyltransferase BF2170. A total of 170 medical drugs containing at least one carboxyl moiety (20 µM) were individually tested against the BF2170-expressing *E. coli* strain under an anaerobic condition. LC–MS analysis identified products of carboxyl methylation showing a mass difference matching +14.016 Da and an increase in RT. **b**, Uniform Manifold Approximation and Projection (UMAP) of the selected carboxyl drugs mapped onto 12,104 small-molecule drugs from DrugBank^28^. Number of drugs in each group is indicated in parentheses. **c**, Chemical and pharmacological classification of the 170 selected drugs and 15 methylation hits. Inner ring, chemical superclass; middle ring, ATC anatomical main group; outer ring, mean LC–MS peak area of methylation product at 24 h of incubation, calculated from 2 independent assay replicates (Supplementary Table 20). **d,** Experimentally confirmed chemical structures of the identified carboxyl-methyl ester prodrugs produced by BF2170. See Supplementary Data for structural validation by LC–MS/MS and comparison with synthetic standards.

## Discussion

For over a century, the prodrug concept has been employed in drug development (*e.g.*, aspirin synthesis), with most prodrugs designed according to medicinal chemistry principles and their bioactivation relying on host enzymes^5,16^. We uncovered a natural microbial route that follows the same chemical principle, showing that the gut microbiota produces chemically diverse drug metabolites with the potential to act as prodrugs. This finding expands our view on prodrug functioning and suggests a novel mechanism by which the gut microbiota can modulate drug pharmacokinetics.

The human gut microbiota has emerged as a central and mechanistically distinct contributor to drug metabolism and disposition. With a growing body of publicly accessible datasets on microbial drug biotransformation (*e.g.*, ChEMBL)^1,3,29^, the field is now poised to shift from descriptive profiling toward mechanistic discovery^30^. Screening-derived metabolomics datasets, which quantify microbial drug biotransformation at scale, offer a reproducible platform to generate and test hypotheses about microbial drug biotransformation, thereby enabling the prioritization of drug metabolites, as well as microbial species and enzymes, for functional interrogation in relevant model systems. Our study illustrates how mining such datasets can functionally link observations of microbial drug biotransformation to pharmacological outcomes. By pinpointing drug metabolites as candidate prodrugs, identifying responsible microbial enzymes, validating their function via intestinal permeability assays, and a gnotobiotic mouse experiment, we uncovered a distinct microbial route of enzymatic prodrug generation *in situ* that enhances systemic drug exposure.

Discovering microbial enzymes involved in drug metabolism presents a major opportunity to gain a mechanistic understanding of microbe–drug interactions and inform drug discovery and development^31^. Although burgeoning computational tools have improved protein annotations^32^ and enabled prediction of enzymes responsible for drug metabolism^31,33^, functional validation of these putative enzymes remains scarce. We demonstrate that combining genetic screening and metagenome mining offers a successful strategy to identify functional drug-modifying enzymes. Applied to *B. fragilis* DSM2151, the genetic GoF screen identified *bf2170* as the sole gene responsible for drug methylation. Notably, this single hit emerged from a pool of thousands of gene candidates including 44 predicted methyltransferase-encoding genes in the strain genome (UniProt query results), underscoring the value of the genome-wide screen for identifying functional genes without gene-by-gene testing. The validated GoF hit then served as a functional anchor to map and experimentally test predicted isofunctional enzymes across 289,232 prokaryotic genomes of the human gut microbiome. This provides a scalable roadmap for expanding the drug–enzyme–metabolite network underlying microbial drug metabolism.

Methyltransferases are widespread in nature and catalyze the methylation of diverse biomolecules across biological systems. Within the gut microbiota, methyltransferases have recently been implicated in physiological functions such as epigenetic regulation and pathogenesis^34,35^. Our study identified a new function of gut bacterial methyltransferases acting on xenobiotics. Drug biotransformation assays revealed carboxyl methylation activity across structurally distinct drug compounds, particularly active on fibrates and profens, underscoring the chemical versatility of these microbial methyltransferases and motivating in-depth investigation into their pharmacological relevance. Future biochemical and structural investigation of these microbial methyltransferases may illuminate the molecular basis of substrate selectivity and support predictive models of microbial drug modifications in the gut. Furthermore, it is likely that the methylation of carboxyl compounds is not restricted to drugs and other xenobiotics, but may also be observed for physiological compounds such as metabolites derived from food, other gut microbes or the host, impacting microbiota–host metabolic interactions.

## Methods

### Experimental materials

Chemicals (Supplementary Table 21), miscellaneous materials and instruments (Supplementary Table 22), plasmids and primers (Supplementary Table 23), and bacterial strains (Supplementary Table 24) used in this study are listed in the Supplementary Information. The identity of all bacterial strains was verified by 16S rRNA gene sequencing (primers 1 and 2) prior to experiments.

### Mining microbial prodrug candidates from the reported metabolomics dataset

The previously reported metabolomics dataset of gut microbial drug metabolism (ChEMBL ID: CHEMBL5303573; MetaboLights accession number: MTBLS896)^1^ was used for mining microbial metabolites as prodrug candidates. Of the 76 gut bacterial strains tested in the original study, 3 strains with incomplete biotransformation data were excluded, leaving 73 strains analyzed and reported in this study. To identify conjugation metabolites, 871 putative drug metabolites were annotated by matching mass differences with respect to parent drugs to common phase II drug biotransformation reactions^17^ with a mass error < 0.002 Da. logD was modelled by the ‘Partitioning bundle’ of Chemaxon Calculators and Predictors (http://chemaxon.com/calculators-and-predictors).

### Genetic gain-of-function screen of *B. fragilis* DSM2151

#### Library preparation

Heterologous expression of the GoF library of *B. fragilis* DSM2151 was prepared as previously described^1^ with following adjustments. The expression vector pZE21^37^ (plasmid 1) was linearized by PCR (primers 3 and 4) and dephosphorylated with calf intestinal alkaline phosphatase (Invitrogen). Genomic DNA from *B. fragilis* DSM2151 was sheared using a g-TUBE (Covaris) set to a target size at 6 kb. Gel-purified DNA fragments (2–10 kb) were ligated with PCR-linearized pZE21 via blunt-end ligation (Thermo Scientific). Gel-purified ligation products were electroporated into *E. cloni* 10G (Lucigen), grown on Luria Broth (LB) supplemented with kanamycin (50 µg/ml) (LB-Kan) agar plates at room temperature for 3 days to form single colonies, and arrayed into 384-well plates containing 70 μl LB-Kan broth using an automated colony-picking station (Hudson Lab Automation). To assess library quality, 24 colonies were randomly picked for PCR amplification of multiple cloning sites (primers 5 and 6). The resulting PCR products were analyzed by gel electrophoresis, confirming high cloning efficiency (Supplementary Fig. 2a), and Sanger sequenced to estimate the average insert size and to validate unbiased distribution of genomic insert locations (Supplementary Fig. 2b). Overnight-grown, arrayed plates were replicated onto LB-Kan agar plates using a Singer Rotor HDA pinning robot (Singer Instruments) for hit screening, and original arrayed plates were preserved at −70 °C for localizing active clones within hit plates as part of the screening protocol (see below).

#### Library screening

GoF library screening was performed as previously described^1^. All 384 colonies from each replicate library agar plate were collected en masse by scraping and pooled in 750 μl 1/2-diluted gut microbiota medium (GMM)^38^. The cell suspension (180 μl) was mixed with 1/2-diluted GMM containing BEZ (20 μl; final 1.5 μM) and incubated anaerobically at 37 °C in a 96-well plate. Aliquots (20 μl) were collected at 0, 1.5, 3, 6, 12, and 24 h and analyzed by LC–MS to identify hit plates (see Supplementary Fig. 2c). To localize methylation-active clones, hit plates showing BEZ-methylating activity were defrosted and replicated into 384-well LB-Kan plates. Overnight-grown cultures were row- and column-pooled using a liquid-handling platform (Agilent Bravo), centrifuged, and resuspended in 200 μl 1/2-diluted GMM containing BEZ to test for methylation (Supplementary Fig. 2d for pooling procedure). One active hit clone (library index P56-D2) was colony-purified and 4 independent colonies per clone were re-tested for BEZ-methylating activity, and 2 verified clonal cultures were selected for Sanger sequencing of inserts (primers 5 and 6).

#### Hit validation by target gene cloning

Two genes, *bf2169* and *bf2170*, identified in the methylation-active clone were validated individually by heterologous expression. PCR-amplified open reading frames including flanking regions (primers 7–10) were cloned into pZE21 plasmids via Gibson assembly (NEBuilder HiFi DNA Assembly), transformed into *E. cloni*, and tested for BEZ-methylating activity under anaerobic conditions.

### Genetic manipulation of *Bacteroides fragilis*

#### Gene deletion of bf2170

An in-frame, unmarked deletion of *bf2170* (also annotated as BF9343_2083; UniProt accession number: A0A380YZ71) was generated in *B. fragilis* DSM2151 using an allelic exchange protocol with the counter-selectable suicide plasmid pLGB13^39^ (plasmid 2), delivered via a diaminopimelic acid (DAP)-auxotrophic *E. coli* dual conjugation donor/cloning strain (DATC) strain^40,41^. A 2-kb insert lacking *bf2170*, consisting of 1-kb flanking regions upstream and downstream of the open reading frame, was assembled by overlap extension PCR^42^ (primers 11–14), and integrated into pLGB13 via restriction sites EcoRV and PstI. The construct was PCR-verified (primers 15 and 16), transformed into the electrocompetent donor *E. coli* DATC, and conjugated into *B. fragilis*. For conjugation, overnight dense cultures of the *E. coli* donor in LB (supplemented with 100 µg/ml of ampicillin and 0.3 mM of DAP) and the *B. fragilis* recipient in modified Gifu Anaerobic Broth (mGAM) were mixed (1 ml + 1 ml) and spun down (12,000 × g, 2 min). Pellets were resuspended in 100 μl of mGAM with DAP (0.3 mM) and spotted onto a prewarmed mGAM agar plate without DAP and incubated aerobically at 37 °C for 20 h. Cells from the mating spot were plated on an mGAM agar plate supplemented with erythromycin (5 μg/ml) to grow anaerobically at 37 °C. After 36 h, single colonies were picked and re-streaked for isolation. To resolve co-integrates, isolated clones were plated on mGAM agar plates with anhydrotetracycline (200 ng/ml) and incubated anaerobically overnight. Single colonies were purified by re-streaking and screened by PCR (primers 17 and 18) to confirm the *bf2170* deletion.

#### Complementation of bf2170

Gene complementation at various expression levels was performed using pNBU2-derived plasmids (plasmids 3−9) as previously described^36^. PCR-amplified *bf2170* with flanking regions (primers 19 and 20) was cloned into PCR-linearized pNBU2 plasmids (primers 21 and 22) via Gibson assembly, and transformed into the electrocompetent *E. coli* DATC donor strain, and conjugated into the Δ*bf2170* mutant strain using the same protocol described above. Complementation using the strongest promoter (P1E6) was unsuccessful, likely due to enzyme toxicity at a high expression level.

### BF2170 homolog analysis

#### Searching BF2170 homologs in the reported 73-strain panel

To identify sequence homologs of BF2170 in the reported 73-strain panel^1^, NCBI BLASTP+ (v2.16.0) was employed with thresholds of >25% amino acid sequence identity and E-value < 10^-3^ as permissive criteria to collect all putative homologs. The amino-acid sequence of BF2170 served as the query. Strains lacking publicly available genomes (4) were excluded, yielding a final list of 69 strains for analysis. Strains encoding a putative BF2170 homolog were plotted by their BEZ-methylating activity versus the sequence identity of the respective homolog to examine the relationship between these two parameters.

#### Searching isofunctional proteins of BF2170 in the UHGG collection

The amino acid sequence of BF2170 was queried against the Unified Human Gastrointestinal Protein (UHGP) catalog clustered at 100% amino-acid identity (UHGP-100 v2.0.2)^24^, downloaded from https://ftp.ebi.ac.uk/pub/databases/metagenomics/mgnify_genomes/. Command-line NCBI BLASTP+ (v2.16.0) was employed to search proteins with high sequence similarity to BF2170 using thresholds of ≥50% amino acid sequence identity, ≥50% query coverage, and E-value < 10^-6^. The resulting hit proteins were called ‘putative isofunctional proteins of BF2170’ throughout the study. Nine putative isofunctional proteins of BF2170 from 8 commercially available strains (Supplementary Table 11) were validated for BEZ-methylating activity through cloning (primers 23–38) and heterologous expression in *E. coli* followed by *in vitro* assays (see ‘*Hit validation by target gene cloning*’). Clustal Omega (v1.2.4)^43^ was used for multiple sequence alignment of BF2170 and 9 additionally tested isofunctional proteins of BF2170 (job ID: clustalo-I20250728-144355-0682-46598267-p1m; see Supplementary Method for protein structure prediction using Boltz-1 (v0.4.1)^44^). To visualize the phylogenetic distribution of putative isofunctional proteins of BF2170 within UHGG (v2.0.2), bacterial phylogeny was reconstructed using the bact120 protein alignment file downloaded from the FTP server of MGnify^45^. IQ-TREE (v2.3.3)^46^ was used to generate a maximum-likelihood phylogenetic tree for the 4,716 representative bacterial species. The phylogenetic tree was midpoint-rooted and visualized in iTOL (v7.2.1)^47^ (see Supplementary Method for parameters to construct phylogenetic trees).

### *In vitro* drug biotransformation assay

#### Biotransformation assays using bacterial cultures

Frozen glycerol stocks (−70 °C) of gut bacterial strains or *E. cloni* clones expressing methyltransferases were streaked onto brain–heart infusion supplemented with 10% defibrinated horse blood (BHI blood) agar plates and incubated anaerobically at 37 °C for 24–48 h. Single colonies were then inoculated into 4 ml pre-reduced mGAM medium and cultured anaerobically for 24 h to generate precultures. The assay was initiated by adding precultures at 10% volume into 2-fold diluted, pre-reduced mGAM containing BEZ (final 2 µM) with a final volume of 200 µl, mixed directly in a 96-well plate (Thermo Scientific 267544). At 0, 1.5, 3, 6, 12, and 24 h of anaerobic incubation at 37 °C, 20 µl aliquots were collected in V-bottom 96-well polypropylene plates (Abgene 10304513), and snap-frozen. Samples were stored at −70 °C prior to organic solvent extraction (see *‘Sample preparation for LC–MS analysis’*) and LC–MS analysis.

#### Carboxyl drug methylation screen

A total of 170 medical drugs containing aliphatic carboxylic acid moiety were selected from a customized in-house drug library (Selleck Chemicals; see Supplementary Table 20 for drug information). The selected drugs and 12,104 small-molecule drugs from DrugBank^28^ (v5.1.13; drug entities were filtered with molecular weight of <1,210 Da and with available SMILES) were used to generate Morgan fingerprints using the RDKit (v2025.9.1; rdFingerprintGenerator.GetMorganGenerator function with radius of 2 and fpSize of 2048), which were used for UMAP visualization using Jaccard distance (umap-learnpackage in Python, v0.5.9). Chemical taxonomy was assigned via ClassyFire^48^ using the ChemOnt ontology (v2.1; http://classyfire.wishartlab.com/downloads). ATC codes were retrieved from DrugBank and manually curated. The BF2170-expressing *E. coli* strain (as generated above in ‘*Hit validation by target gene cloning*’), or a control strain with an empty pZE21 vector, was incubated against each drug (20 µM in 1/2-diluted GMM) following the biotransformation protocol as described above, with samples collected at 0, 12, and 24 h of anaerobic incubation at 37 °C. To increase throughput, extracted samples were pooled by column (*i.e.*, each pool combined samples from 8 wells in the same column of a 96-well plate) and analyzed by LC–MS. Identified methylated products (see ‘*Data analysis*’ in ‘*LC–MS methods and data analysis*’) were validated by LC–MS/MS comparison with synthetic standards (see Supplementary Method for chemical synthesis of drug carboxyl-methyl esters and Supplementary Data for LC–MS/MS spectra)

### Caco-2 transwell assay to assess trans-epithelial permeability

Caco-2 transwell assays were performed as previously described^49^. In brief, Caco-2 cells seeded at a density of 100,000 cells/well in 24-well transwell inserts (pore size of 0.4 µm; Millicell) in standard 24-well plates were cultured for 21 days to form polarized monolayers. To assess the apical-to-basolateral trans-epithelial permeability, 500 µl of Hank’s balanced salt solution with calcium and magnesium at pH 7.4 (HBSS) containing BEZ or BEZ-Me (40 µM; low-dose assay, 2 µM) and 7 marker compounds indicative of transport functionality (antipyrine, haloperidol, propranolol, and warfarin as permeable markers; terbutaline, etoposide, and nadolol as non-permeable markers; 1 µM each) were applied to the apical side, whereas the basolateral side was filled with 500 µl of HBSS. To assess the basolateral-to-apical permeability, BEZ or BEZ-Me was added to the basolateral side, while the marker compounds were added to the apical side. Samples were collected over 4 h by collecting 10 µl aliquots (at 0, 10, 20, 30, 40, 60, 90, 120, 150, 180, 210, and 240 min) from both sides of the transwell setups, and stored at −70 °C prior to extraction and LC–MS analysis. Functional integrity of the barrier was validated by concentration kinetics of marker compounds and cell viability was assessed after the assays using CellTiter-Blue (Promega) according to the manufacturer’s protocol and as previously described^49^. The apparent permeability coefficient (P_app_) of BEZ was calculated as P_app_ = (dC_BEZ_/dt × V_r_) / (A × C_0_), where dC_BEZ_/dt is the slope of the BEZ concentration−time curve in the receiver side, V_r_ is the initial volume of the receiver side, C_0_ is the initial concentration of BEZ or BEZ-Me in the donor side, and A is the area of the transwell membrane. For experiments in which BEZ-Me was applied, P_app_ was calculated using data from the time window during which BEZ-Me hydrolysis was occurring. Detailed calculation of P_app_ is provided in Supplementary Table 15.

### Gnotobiotic mouse model to assess pharmacokinetics

All mouse experiments were performed using protocols approved by the European Molecular Biology Laboratory (EMBL) Institutional Animal Care and Use Committee (license number 21-002_HD_MZ). Germ-free C57BL/6 mice (purchased from Taconic) were maintained and bred in flexible plastic gnotobiotic isolators (Class Biologically Clean) with a 12-h light/dark cycle and provided a standard, autoclaved mouse chow (5013 LabDiet, Altromin) *ad libitum*. Germ-free status of isolators was routinely monitored by culture-based methods and 16S rRNA PCR (primers 1 and 2). Germ-free status of each animal was confirmed by culture-based methods before experiments.

A total of 53 individually caged GF mice (8−20 weeks old) of mixed sex were used to establish a gnotobiotic mouse model (see Supplementary Table 17 and ‘*Statistics and reproducibility’* for group and batch details). Mono-colonization with *B. fragilis* WT or Δ*bf2170* mutant strains was established by oral gavage of 200 µl of overnight dense bacterial culture with approximately 4 × 10^8^ colony-forming unit (CFU). Fresh fecal materials from the 4^th^ day post colonization were used to determine bacterial loads based on anaerobic CFU, validated using *bf2170*-specific PCR (primers 17 and 18), and LAO checked for contamination through plating and incubation under aerobic conditions. On the 7^th^ day post colonization, mice were orally administered a freshly prepared BEZ suspension solution (5 mg/ml in phosphate-buffered saline) at a single dose of 50 mg/kg of body weight^50^. At 3, 5, 7, or 9 h post drug treatment, mice were euthanized in a CO_2_ chamber. Homogenates of cecum contents were collected in 2.0-ml screw-top tubes with O-ring cap (Merck) and snap-frozen by liquid nitrogen. Blood samples were collected using 0.5-ml lithium heparin tubes (Greiner) by heart puncture post euthanization and centrifuged (2,500 × g at 4 °C, 10 min) to obtain plasma from supernatants. Samples were stored at −70 °C prior to extraction and LC–MS analysis.

### Sample preparation for LC–MS analysis

#### Liquid samples from in vitro assays and mouse plasma

A liquid-handling platform (Agilent Bravo) was used for compound extraction of liquid samples. Sample aliquots were added with 1/4 volume of an internal standard (IS) solution (caffeine-d_9_, diclofenac-d_4_, nafcillin-d_5_, oxfendazole-d_3_, phenylalanine-d_5_, tolfenamic acid-d_4_, tryptophan-d_5_, and warfarin-d_5_, each at 20 µM in water) and 5 volumes of cold organic solvent (acetonitrile:methanol = 1:1, v:v) relative to the aliquot, resulting in a final composition of acetonitrile:methanol:water = 2:2:1 (v:v:v). After incubation at −20°C for at least 1 hour to precipitate proteins, extracts were centrifuged (4,347 × g at 4°C, 10 min), and supernatants were diluted with water (1:1, v:v) for LC–MS analysis. For mouse plasma, a modified IS solution containing BEZ-d_6_, BEZ-Me-d_6_, and the above IS compounds (20 µM each) was used, following the same extraction procedure. Calibration curves for absolute quantification of BEZ and BEZ-Me were prepared by a 5-fold serial dilution (3.2 nM–50 µM in water) processed together with liquid samples within the same batch.

#### Mouse cecum

Pre-weighed thawed cecum homogenate (30−45 mg) was mixed with 0.1 mm zirconia/silica beads (∼150 μl; BioSpec Products) and cold organic solvent (acetonitrile:methanol = 1:1, v:v; 300 µl) containing IS compounds (BEZ-d_6_, BEZ-Me-d_6_, and the other IS compounds as described above, each at 0.8 µM) in a 2-ml screw-top tube (Merck). Samples were homogenized by mechanical disruption with a bead beater (BioSpec Products) set for 2 min on ‘high’ setting at room temperature. After incubation for at least 1 h at –20 °C, samples were centrifuged (12,000 × g at 4 °C, 2 min), and supernatants were diluted with 4 volumes of water for subsequent analysis by LC–MS. Calibration curves for absolute quantification of BEZ and BEZ-Me were prepared using 5-fold serial dilutions (3.2 nM–50 µM in water) spiked with the IS compounds at the same concentrations as in cecum samples and processed together with cecum samples within the same batch.

### LC–MS methods and data analysis

#### Methods for in vitro assay samples

An Agilent QTOF 6546 mass spectrometer equipped with an ESI source coupled to an Agilent 1290 Infinity II UHPLC system was used to analyze samples from *in vitro* biotransformation assays and Caco-2 experiments. Agilent 6550 QTOF mass spectrometer equipped with the same configuration was used to analyze samples from the carboxyl-drug screen. Chromatography was performed with an InfinityLab Poroshell 120 HPH-C18 column (2.1 × 100 mm, 1.9 µm) maintained at 45 °C. Sample extracts (5 µl) were injected and separated using a linear gradient with solvent A (water + 0.1% formic acid) and solvent B (methanol + 0.1% formic acid), ramped from 5% to 95% B over 5.5 min and held at 95% B for 1 min at a flow rate of 0.6 ml/min. Data were acquired in positive-ion mode using a centroid spectral format over *m/z* 100–1500. Online mass calibration was performed using a reference solution containing purine (*m/z* 121.0509) and hexakis(1H,1H,3H-perfluoropropoxy)phosphazene (*m/z* 922.0098), introduced via the secondary ESI source at a constant flow rate of 15 µl/min. For LC–MS/MS analysis, the targeted MS/MS mode was employed with an isolation width set to ‘narrow (∼1.3 *m/z*)’ and a collision energy set to 10, 20, or 40 eV in separate acquisitions.

#### Methods for mouse samples

An Agilent 6495D Triple Quadrupole mass spectrometer coupled to Agilent 1290 Infinity II UHPLC system (LC–QqQ) was used to quantify BEZ and BEZ-Me in mouse samples using the same chromatographic method as described above. Data was acquired in dynamic multiple reaction monitoring (dMRM) mode using quantifier and qualifier ion transitions listed in Supplementary Table 25.

#### Data analysis

For quantifying BEZ and BEZ-Me in *in vitro* assays, LC–QTOF peak areas were adjusted by sample-wise variation of IS intensities within each batch, and converted to absolute concentrations using calibration curves generated from serially diluted standards. Calibration curves were constructed from log_10_-transformed data fitted with a linear model (stats::lm function in R; equation: lm(log_10_(concentration) ∼ log_10_(normalized peak area)), yielding a linear range of 3.2 nM–50 µM (R² > 0.99) and LOQ of 3.2 nM for both compounds.

For the carboxyl-drug screen, methylated products were identified based on the following criteria: (i) a mass difference of +14.016 Da (mass error < 5 ppm) with a matched isotope pattern; (ii) an increase in RT of <15% relative to the parent drug (ΔRT_COO-Me_ > 0 and (ΔRT_COO-Me_/RT_drug_) < 0.15); (iii) an increase in peak area over incubation (Area_0h_ ≤ Area_12h_ < Area_24h_); and (iv) absence of the product signal in control samples using the *E. coli* clone with an empty vector.

For quantifying BEZ and BEZ-Me in mouse samples, LC–QqQ peak areas were normalized to corresponding isotope-labeled IS compounds (BEZ-d_6_ and BEZ-Me-d_6_, respectively), converted to absolute concentration using calibration curves generated from serially diluted standards, and normalized to the weight of the extracted sample. Calibration curves were constructed as described above, yielding linear ranges of 0.384–6,000 and 0.0768–6,000 nmol/g (R² > 0.99) and LOQ of 384 and 76.8 pmol/g for BEZ and BEZ-Me in cecum, respectively; linear ranges of 0.16–500 and 0.0064–500 µM (R² > 0.99) and LOQ of 0.16 and 0.0064 µM for BEZ and BEZ-Me in plasma, respectively.

### Statistics and reproducibility

All *in vitro* experiments were performed once with indicated assay replication in the figure legends. No statistical methods were used to predetermine sample size. Statistics and LC–MS data analyses were performed in RStudio (v4.3.1). For Caco-2 assays (Fig. 3c and Supplementary Fig. 4), pairwise comparison of P_app_ was performed with two-sided Student’s *t*-test with unequal variance (t.test function). For the gnotobiotic mouse experiment (Fig. 3d–f), mice were randomized before allocation to groups and cages. Four independent batches of mice were used (see Supplementary Tables 17 for the number of mice used per time point of each batch). In the first 3 batches (January−June 2024; *n* = 45), mice were randomly split into two colonization groups with balanced sex distribution and each mouse was sampled at a pharmacokinetic time point at 3, 5, 7, or 9 h post oral administration (*n* = 5−6 per group per time point). Statistical analyses of plasma BEZ kinetics (Fig. 3f) were performed on log_10_-transformed concentrations fitted with a two-way linear model (lm function; log_10_(concentration) ∼ colonization group * time). Pairwise comparisons between colonization groups at each time point were conducted using estimated marginal means (emmeans function; ∼ colonization group | time) and *P* values were calculated using two-sided Tukey’s post hoc tests. To repeat results at the 9-h time point, the 4^th^ batch (March 2025; *n* = 8; 4 mice per colonization group) was further conducted, yielding results consistent with those obtained in 2024. Sensitivity analysis including the 4^th^ batch did not change statistical significance; therefore, all batches were pooled for reporting. The other experiments conducted were not randomized.

## Supplementary Figures

**Supplementary Fig. 1.**
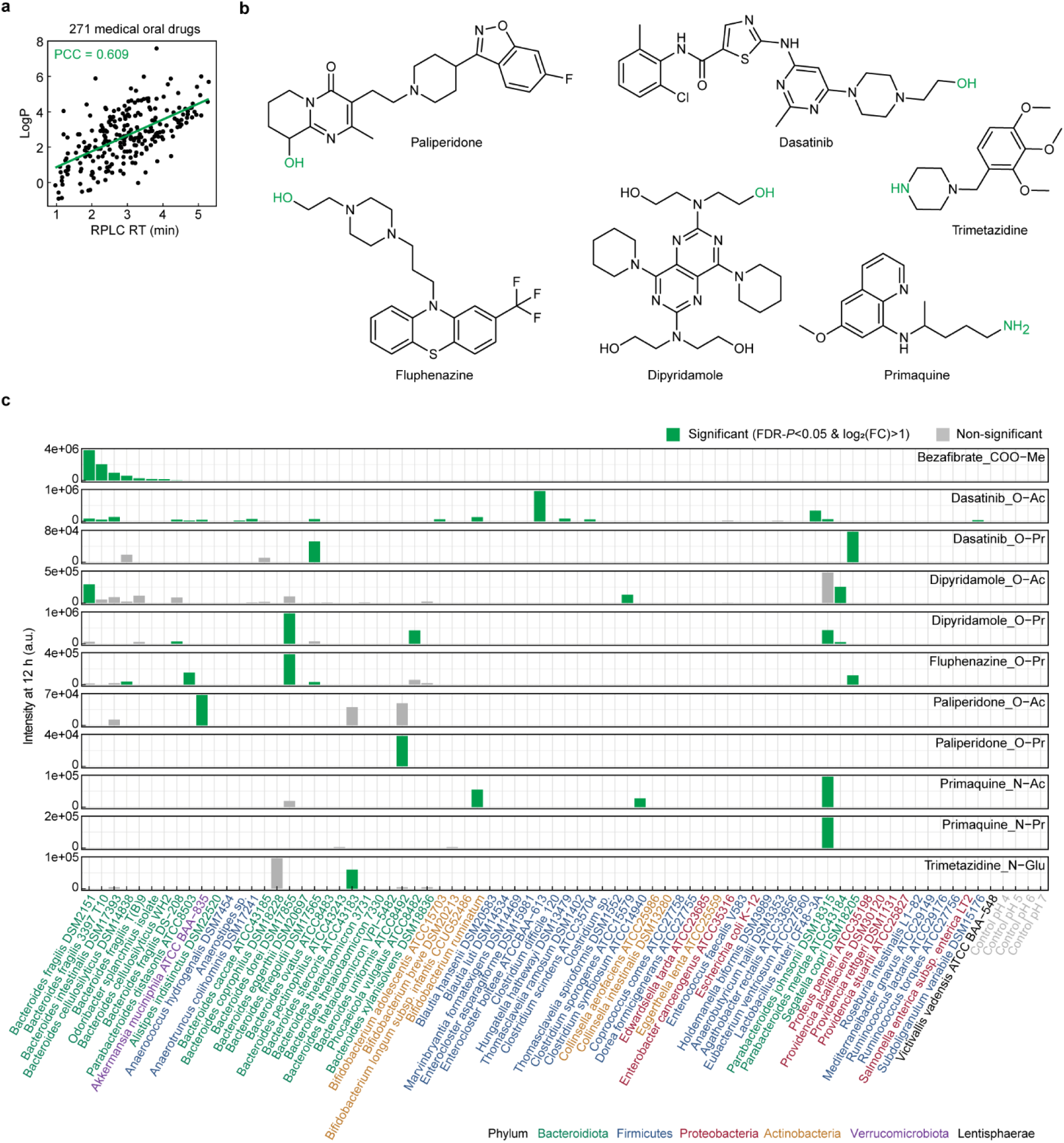
Diverse gut bacteria generate prodrug-like drug metabolites through conjugative biotransformation reactions. **a**, The 271 medical drugs previously tested for gut bacterial biotransformation^1^ show a positive correlation between measured RPLC RT and lipophilicity (logP). Data was fitted with a linear model and linearity was assessed with Pearson correlation coefficient (PCC). **b**, Chemical structures of drugs yielding prodrug-like metabolites. Conjugatable functional groups are highlighted in green. **c**, Reported microbial biotransformation activity producing 11 prodrug-like metabolites^1^. Conjugation form of each drug is indicated in sub-panel titles (‘drug name_conjugation form’). Me, methylation; Ac, acetylation; Pr, propionylation; Glu, glycosylation. Bars represent mean LC–MS peak area calculated from 4 independent cultures. Statistical significance for comparisons between 12 h and 0 h was determined by FDR-adjusted *P* < 0.05 and log_2_(fold change) > 2.

**Supplementary Fig. 2.**
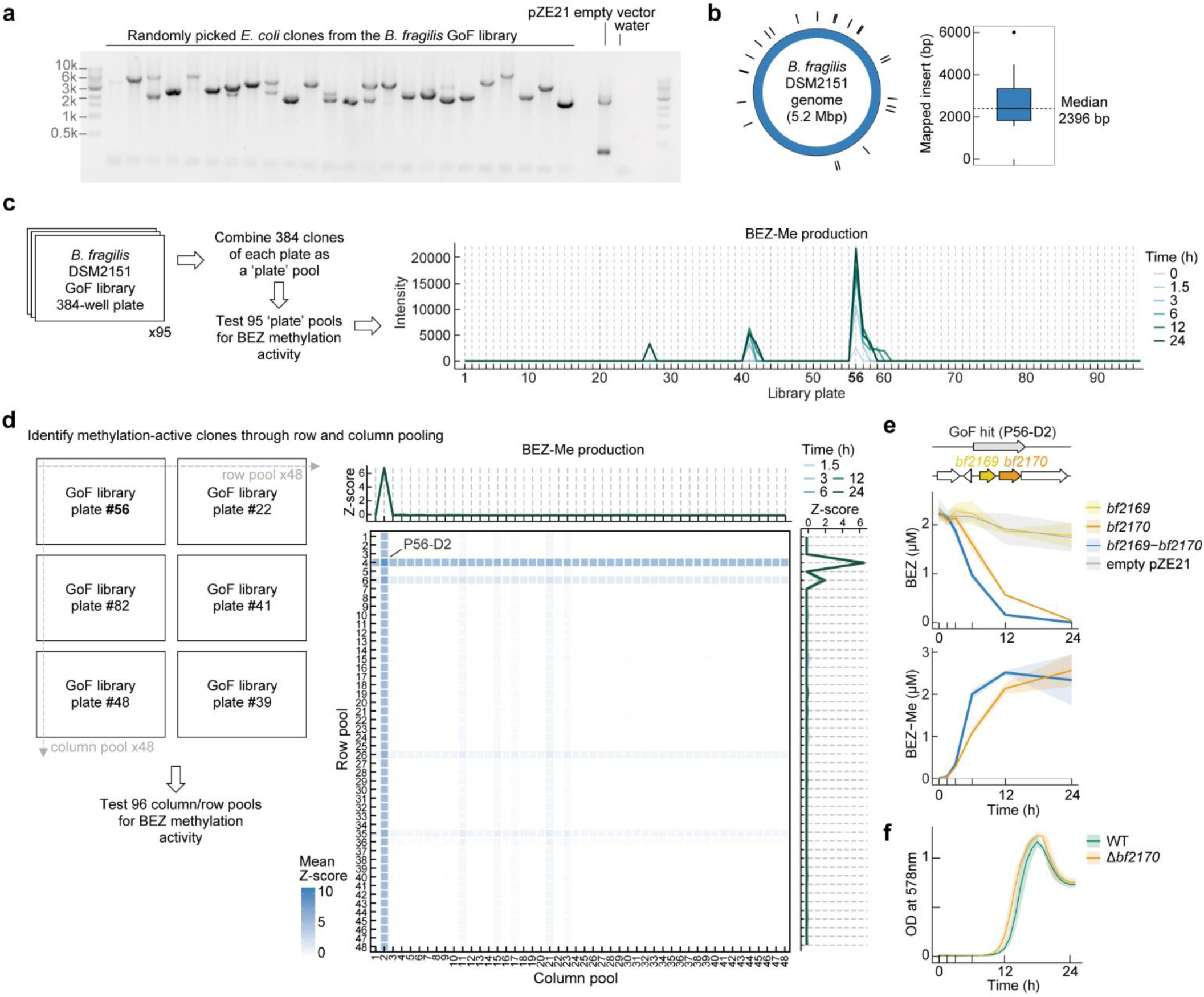
Genetic gain-of-function library of *Bacteroides fragilis* DSM2151: library quality assessment and activity screening identifying *bf2170* responsible for BEZ methylation. **a** and **b**, Quality assessment of the *B. fragilis* DSM2151 GoF library. Transformation efficiency (a) and insert length (b) were assessed by PCR-amplifying multiple cloning sites of 24 randomly picked *E. coli* clones (clone with an empty pZE21 vector as control). Gel image (a) indicates 100% transformation efficiency. Mapping the insert sequences to the strain’s genome indicates an unbiased distribution of inserts and a median insert length of 2,396 bp. **c** and **d**, Experimental procedure for identifying BEZ-methylating clones in the GoF library. All 36,480 GoF clones were pooled in a plate-wise manner and tested for BEZ methylation in anaerobic conditions (c). Subsequently, to locate active clones within the active plate (#56), the active plate #56 alongside 5 otherwise library plates as controls were pooled by rows and columns in a 3×2 configuration, and tested for BEZ-methylating activity (d). Comparing BEZ-methylating activity across 48 column pools and 48 row pools located an active clone at D2 of plate #56 (library index P56-D2). **e**, Open-reading-frame cloning to functionally validate genes covered by the GoF insert region of the hit clone. *E. coli* clones heterologously expressing *bf2169*, *bf2170*, or the combined genomic region from *bf2169* to *bf2170* were tested for BEZ-methylating activity under an anaerobic condition (2 µM of BEZ as substrate). Lines and shaded areas depict mean and s.d., respectively, calculated from 4 independent cultures. **f**, Anaerobic growth curves of *B. fragilis* WT and the Δ*bf2170* mutant strain. Lines and shaded areas depict mean and s.d., respectively, calculated from 5 independent cultures.

**Supplementary Fig. 3.**
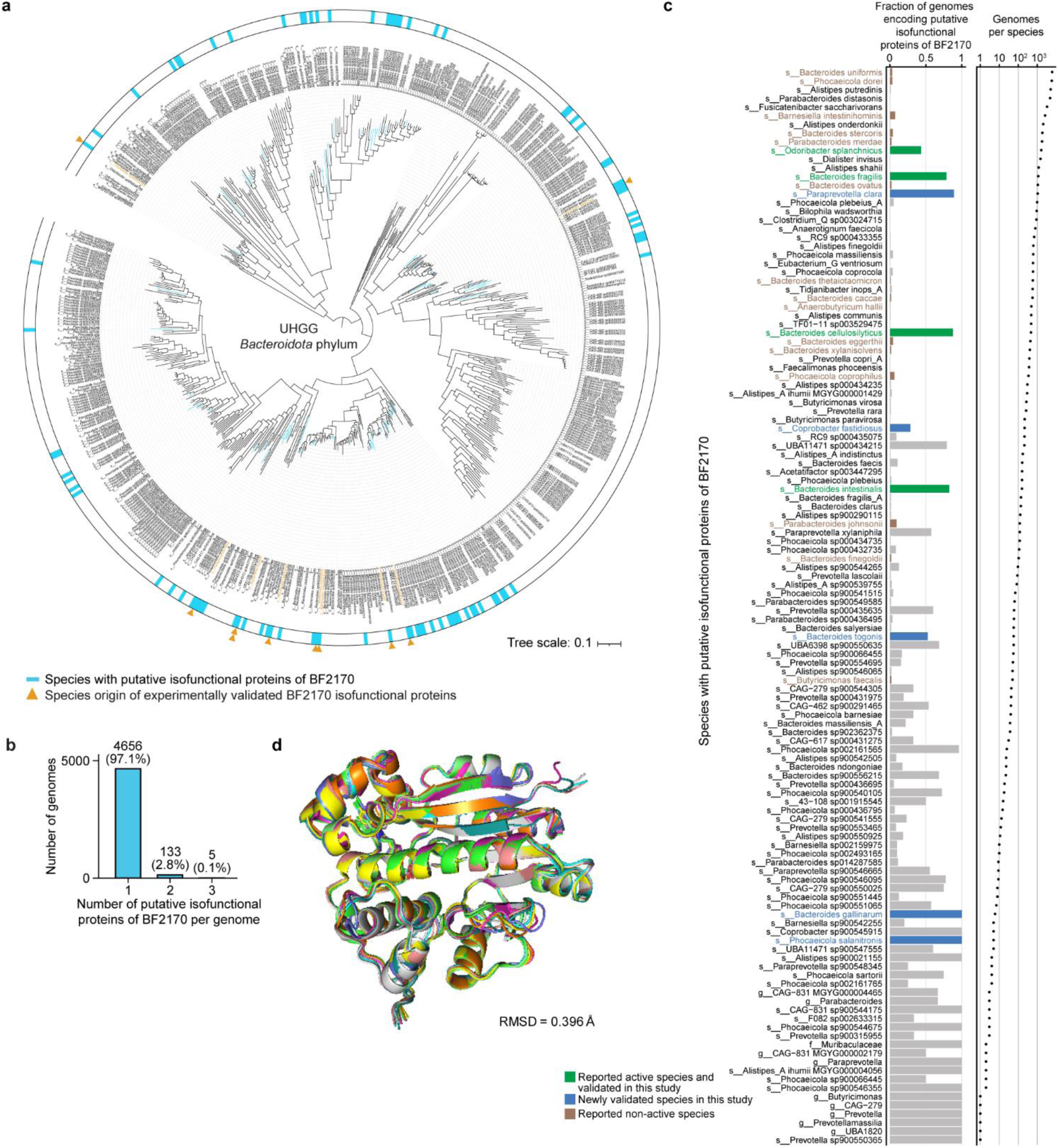
Isofunctional proteins of BF2170 are widely distributed among gut bacteria of the *Bacteroidota* phylum. **a**, The UHGG subtree of the *Bacteroidota* phylum. Light blue bars, 124 species with putative isofunctional proteins of BF2170 (protein sequence identity > 60%); orange triangles, 10 functionally validated proteins originating from 9 species in this study. **b**, Number of putative isofunctional proteins of BF2170 per bacterial genome. **c**, Fraction of genomes encoding putative isofunctional proteins of BF2170 in each hit species. Species from which BF2170 and the 9 validated isofunctional proteins of BF2170 originate are color-coded: green, species with reported BEZ-methylating activity^1^; blue, species newly validated in this study; dark-brown, species reported as non-active^1^. **d**, Predicted structural alignment of BF2170 and the 9 validated isofunctional proteins of BF2170. The predicted BEZ-protein complexes were superimposed and the mean root-mean-square deviation (RMSD) is indicated.

**Supplementary Fig. 4.**
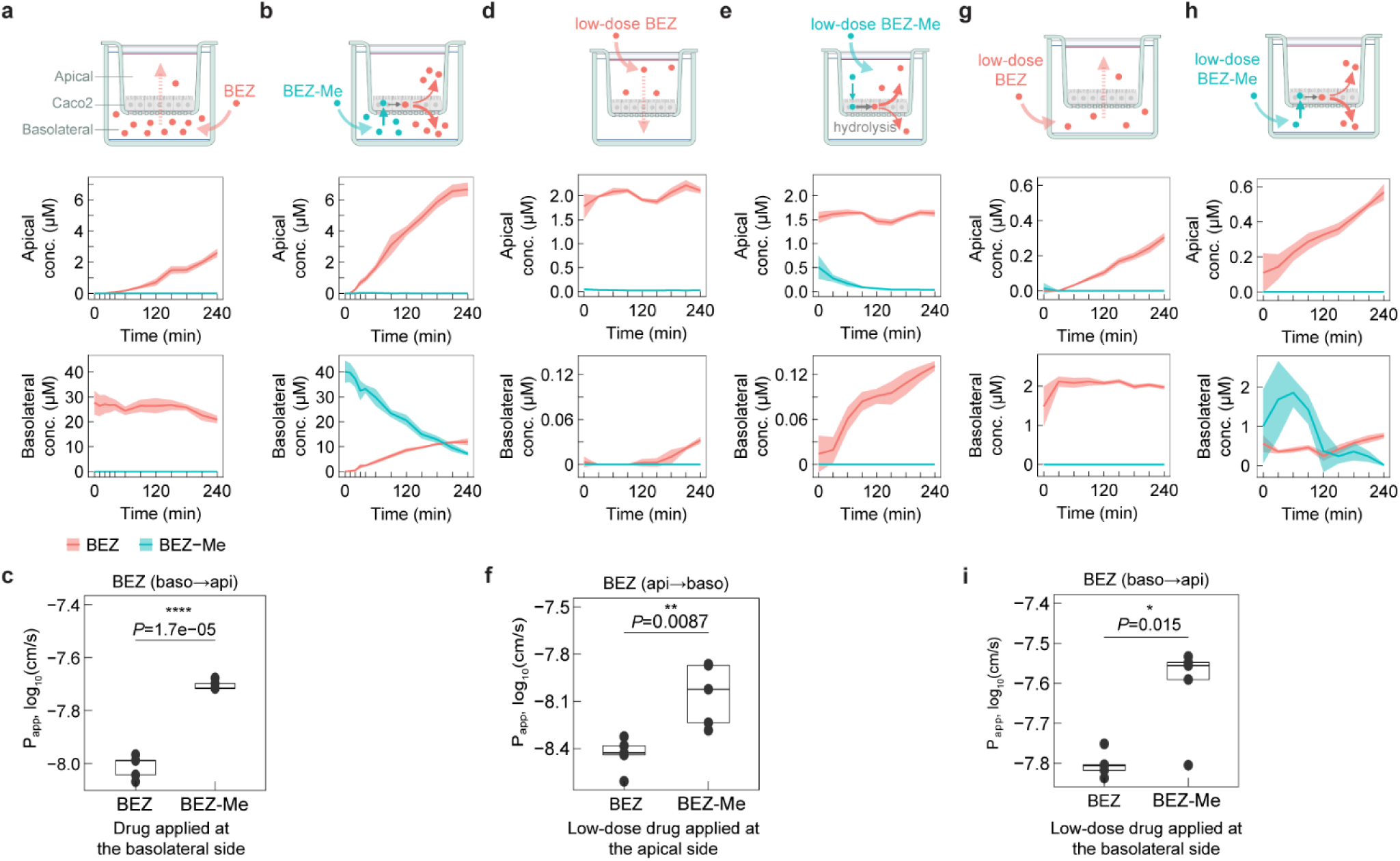
Bidirectional trans-epithelial permeability of BEZ assessed by Caco-2 transwell assays. **a**, **b**, **d**, **e**, **g**, and **h**, Concentration kinetics of BEZ and BEZ-Me when BEZ (a) or BEZ-Me (b) was applied to the basolateral side, and when a low-dose drug (2 µM) was applied to either the apical or basolateral side ((d), BEZ to apical; (e), BEZ-Me to apical; (g), BEZ to basolateral; (h), BEZ-Me to basolateral). Lines and shaded areas depict mean and s.d., respectively, calculated from 5 independent cell assays. **c**, **f**, and **i**, Directional P_app_ of BEZ calculated from each experimental condition ((c) and (i), basolateral-to-apical direction; (f), apical-to-basolateral direction). Pairwise comparison was performed with two-sided Student’s *t*-test with unequal variance. * *P* < 0.05. ** *P* < 0.001. **** *P* < 0.0001. Transwell symbol was created with BioRender.com.

## Data availability

All source data of the figures presented in this study are provided in the Supplementary Tables. Raw mass spectrometry data are deposited in the MetaboLights repository (provisional accession number REQ20251112214584).

## Code availability

Codes for bioinformatics and cheminformatics analyses are available at https://github.com/ZimmermannLab/Prodrugs.

## Acknowledgements

We thank members of the Zimmermann group for discussions and support, especially D.M.Selegato, R.P.Jacoby, and V.Paul for mass spectrometry support, F.Rajer and L.M.Margara for support of molecular biology protocols, E.Mastrorilli for biostatistics discussion and support with data management, M.S.A.Santiago, N.Denisov and K.Fu for support with mouse experiments, M.Beliaeva for discussion on enzyme biochemistry, and C.G.P.Voogdt for support with microbial genetics. We acknowledge B.J.Bartmanski from the Zimmermann-Kogadeeva lab at EMBL and K.Erbstein for assessing the GoF library quality. We acknowledge the Chemical Synthesis Core Facility (CSCF) at EMBL, especially M.Kucukdisli and F.Braun, for support of chemical synthesis. We acknowledge Laboratory Animal Resources team at EMBL, especially C.Grossmann and S.Rudzky, for support of gnotobiotic mouse experiments. We acknowledge the Metabolomics Core Facility at EMBL for support of analytical instruments. We acknowledge Typas lab at EMBL for sharing high-throughput culturomics robots. T.-H.K. was in part supported by the MSCA Postdoctoral Fellowship (101107447) and the Humboldt Research Fellowship for Postdocs. A.S. was in part supported by the MSCA Postdoctoral Fellowship (101109251). This study is supported by the European Research Council (ERC) (GutTransForm-101078353).

## Author information

### Author contribution

T.-H.K. and M.Z. conceptualized the project. T.-H.K. conducted the metabolomics data mining, *in vitro* drug assays, bacterial genetic modifications, and mass spectrometry analyses (unless otherwise specified). M.W.G. and T.-H.K. generated the *B. fragilis* GoF library. A.S. performed the bioinformatics analyses involving the UHGG dataset. R.G.E. performed the homolog analysis in the reported metabolomics dataset, the protein-ligand complex prediction, and the multiple sequence alignment. L.-Y.C. and T.-H.K. carried out the heterologous gene expression and validated the methyltransferases. A.B.-N. and T.-H.K. designed and performed the Caco-2 transwell experiments. G.E.M. and T.-H.K. designed and executed the gnotobiotic mouse experiment. M.Zu. performed the cheminformatics analyses on the carboxyl drug screen. T.-H.K. and M.Z. wrote the manuscript. M.Z. supervised the project, participated in experimental design, and provided scientific guidance and laboratory resources. All authors contributed to manuscript editing and approved the final draft of the manuscript.

## Supplementary information

### Supplementary Tables

This file contains Supplementary Tables 1–25; see the table of contents for details.

### Supplementary Data

High-resolution LC–MS/MS spectra of bacterial carboxyl-methyl drug metabolites and matched synthetic standards.

### Supplementary Methods

Additional methods for anaerobic cultivation, parameters for phylogenetic tree construction, protein structural prediction and alignment, chemical synthesis of carboxyl-methyl ester prodrugs, and detailed instrumental parameters used in high-throughput LC–QTOF and LC–QqQ analyses.

