## Supplementary Data for "Gut bacteria generate prodrugs in situ increasing systemic drug exposure"

for

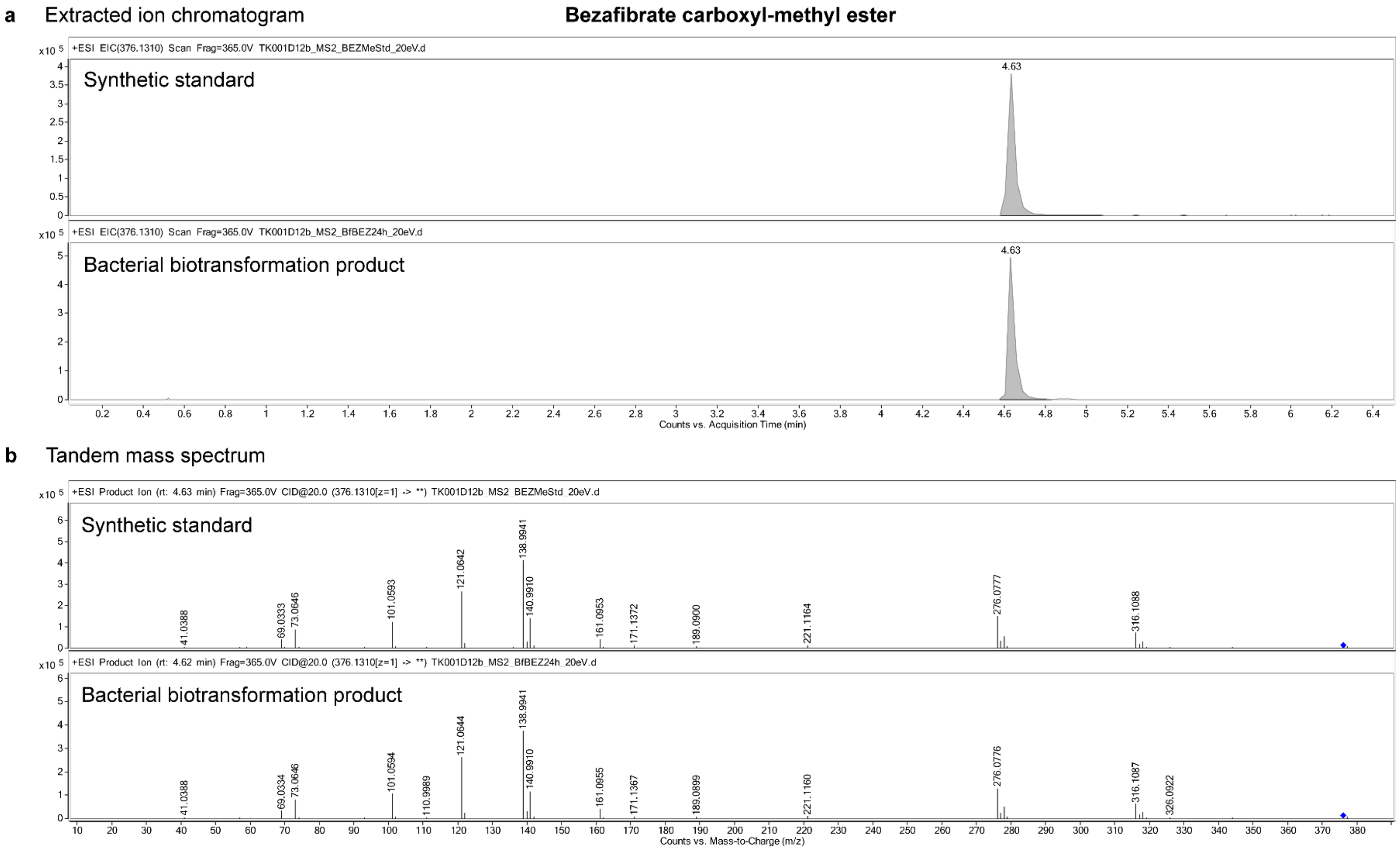

**Figure S1 | LC–MS/MS validation of bezafibrate carboxyl methyl ester produced by bacterial biotransformation.**

**a**, Extracted ion chromatograms and **b**, tandem mass spectra of the bacterial carboxyl-methylation product of bezafibrate and the synthetic standard. Blue diamond, precursor ion.

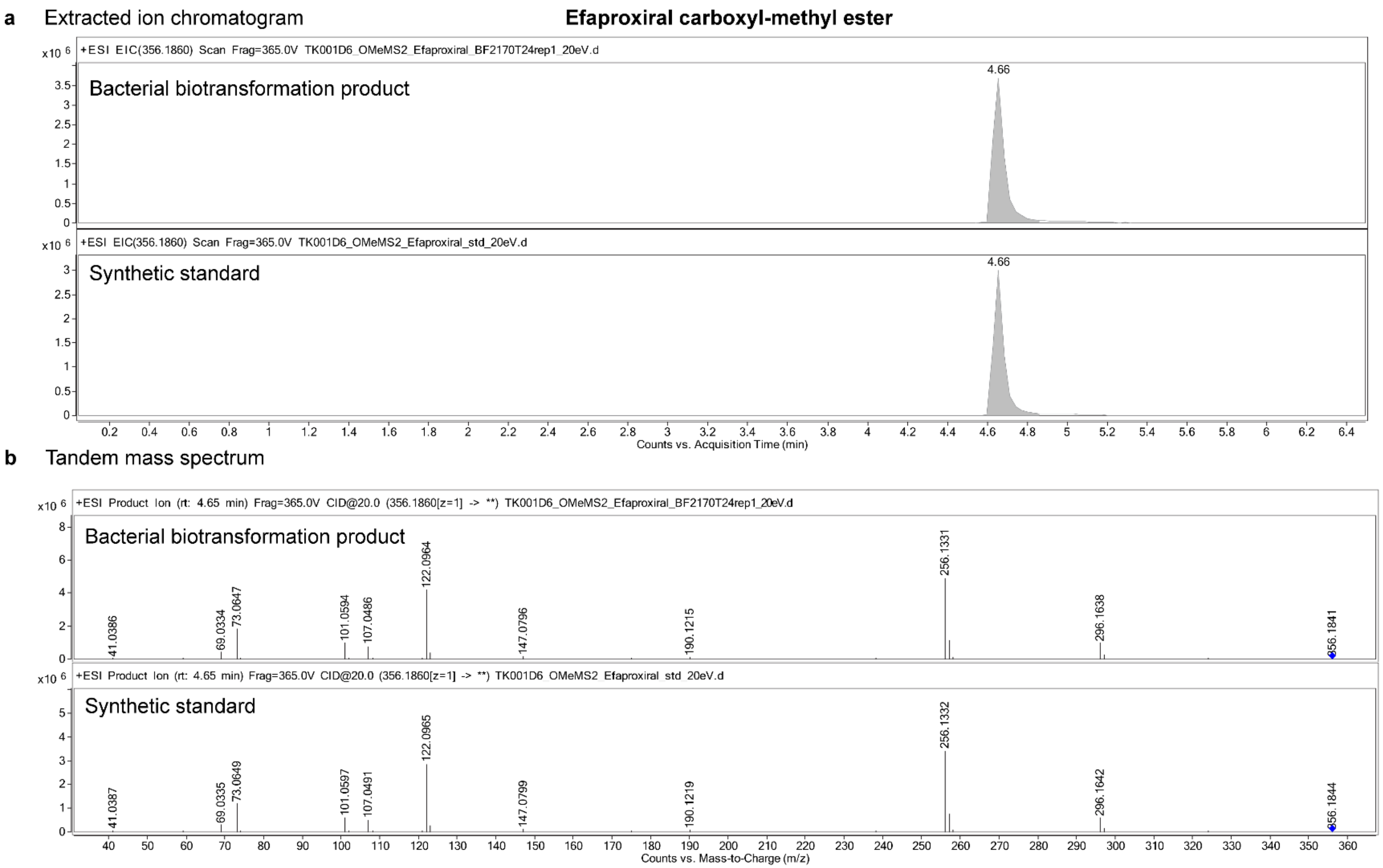

**Figure S2 | LC–MS/MS validation of efaproxiral carboxyl methyl ester produced by bacterial biotransformation.**

**a**, Extracted ion chromatograms and **b**, tandem mass spectra of the bacterial carboxyl-methylation product of efaproxiral and the synthetic standard. Blue diamond, precursor ion.

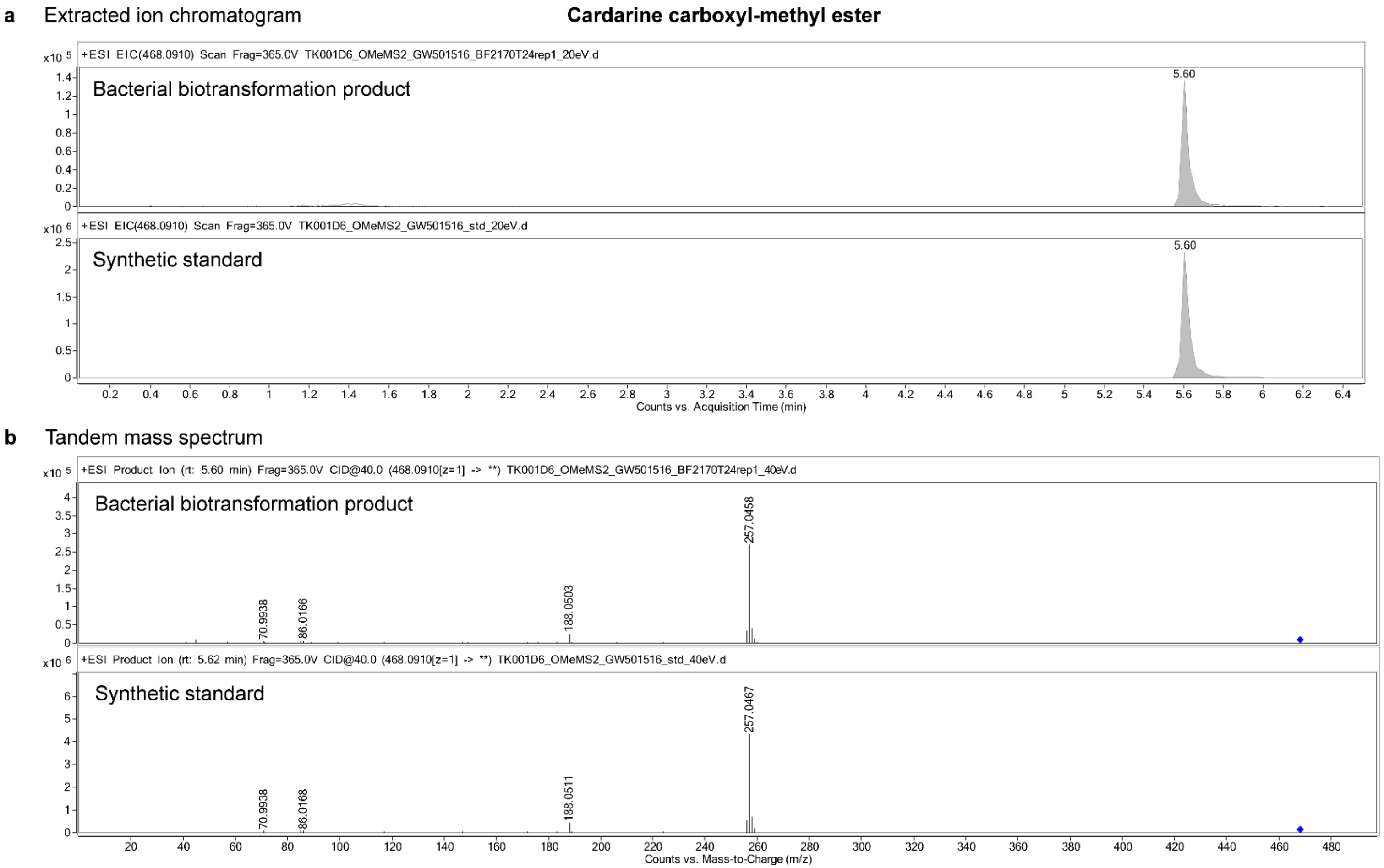

**Figure S3 | LC–MS/MS validation of cardarine carboxyl methyl ester produced by bacterial biotransformation.**

**a**, Extracted ion chromatograms and **b**, tandem mass spectra of the bacterial carboxyl-methylation product of cardarine and the synthetic standard. Blue diamond, precursor ion.

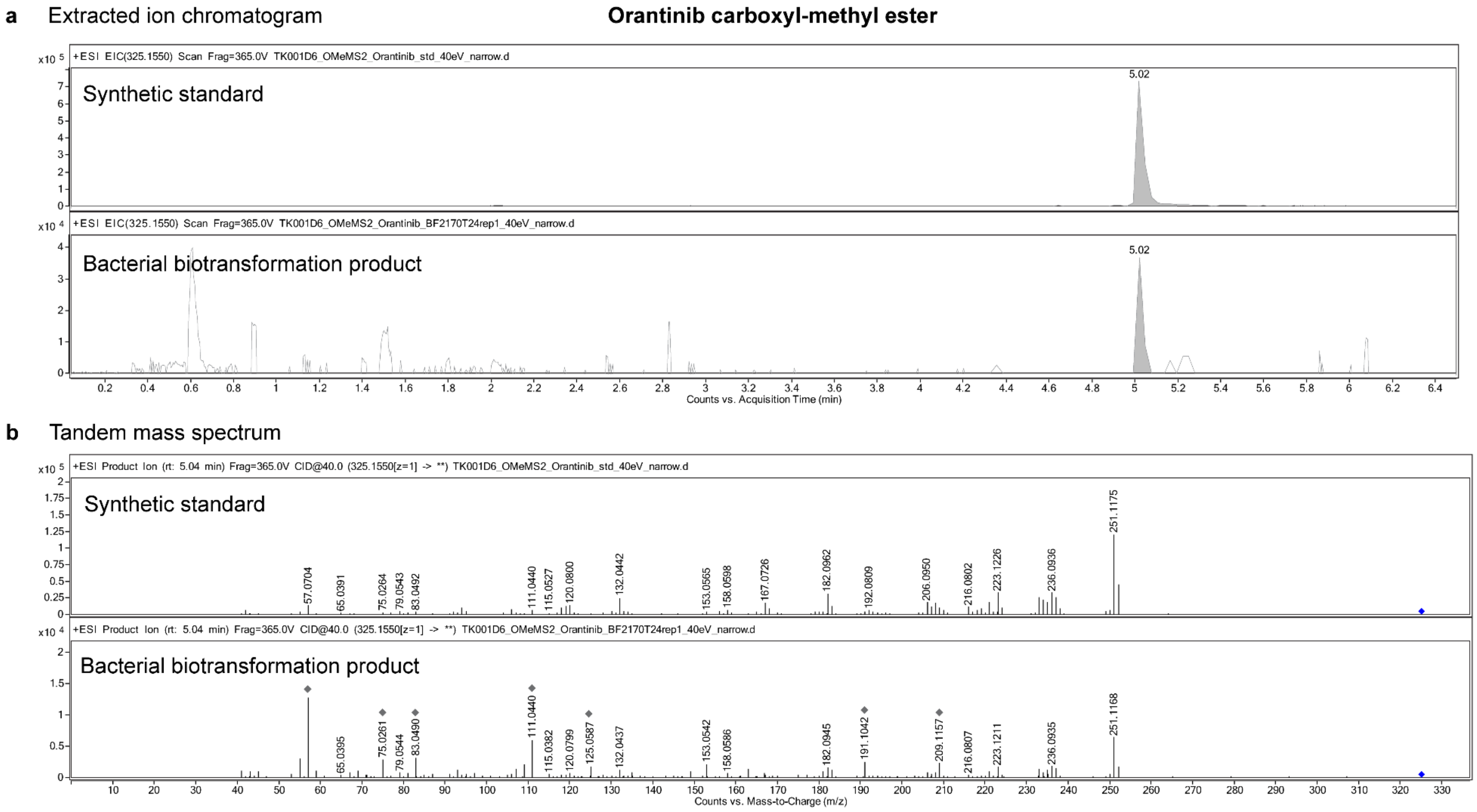

**Figure S4 | LC–MS/MS validation of orantinib carboxyl methyl ester produced by bacterial biotransformation.**

**a**, Extracted ion chromatograms and **b**, tandem mass spectra of the bacterial carboxyl-methylation product of orantinib and the synthetic standard. Blue diamond, precursor ion; grey diamond, background signals.

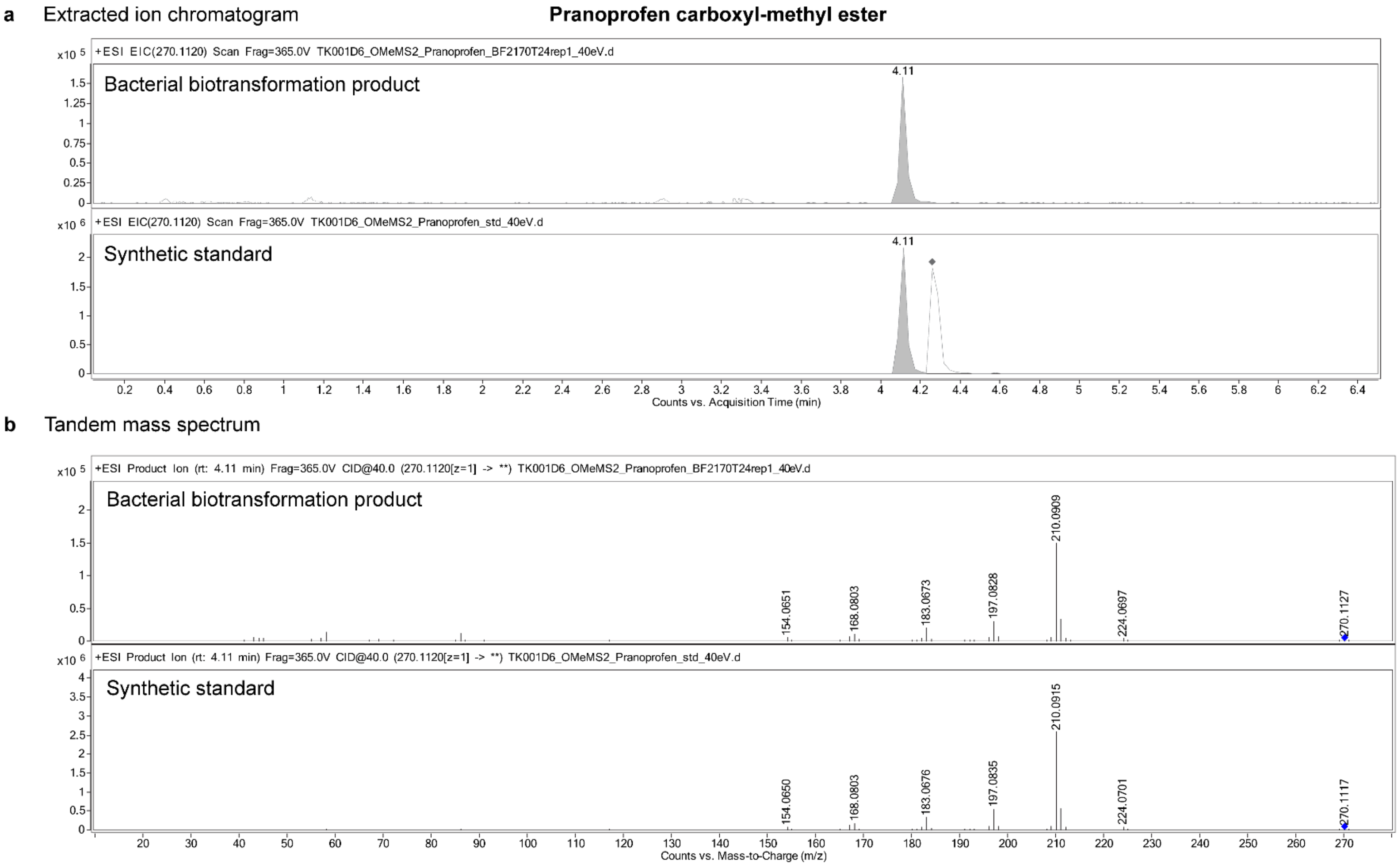

**Figure S5 | LC–MS/MS validation of pranoprofen carboxyl methyl ester produced by bacterial biotransformation.**

**a**, Extracted ion chromatograms and **b**, tandem mass spectra of the bacterial carboxyl-methylation product of pranoprofen and the synthetic standard. Blue diamond, precursor ion; grey diamond, background signals.

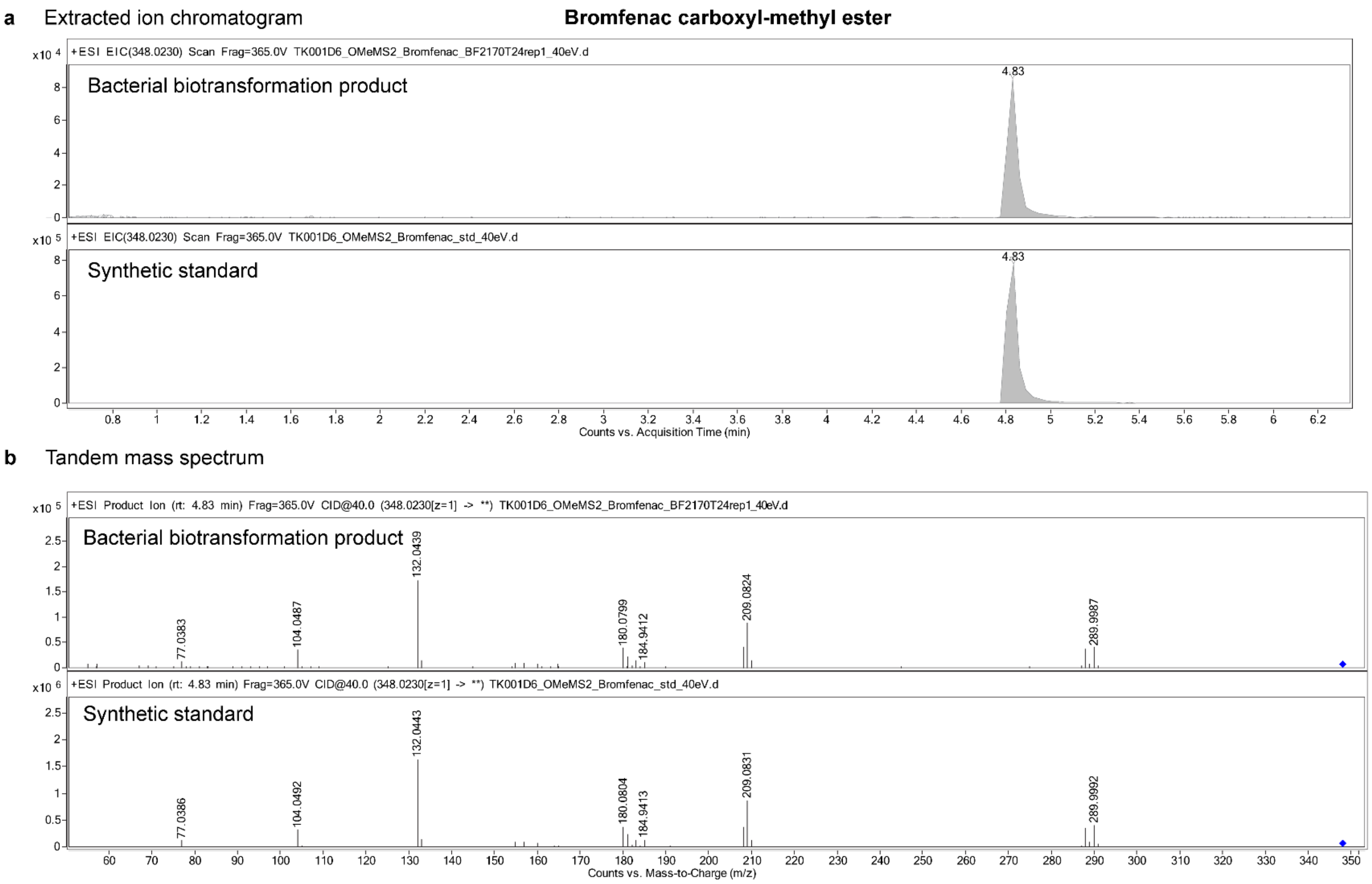

**Figure S6 | LC–MS/MS validation of bromfenac carboxyl methyl ester produced by bacterial biotransformation.**

**a**, Extracted ion chromatograms and **b**, tandem mass spectra of the bacterial carboxyl-methylation product of bromfenac and the synthetic standard. Blue diamond, precursor ion.

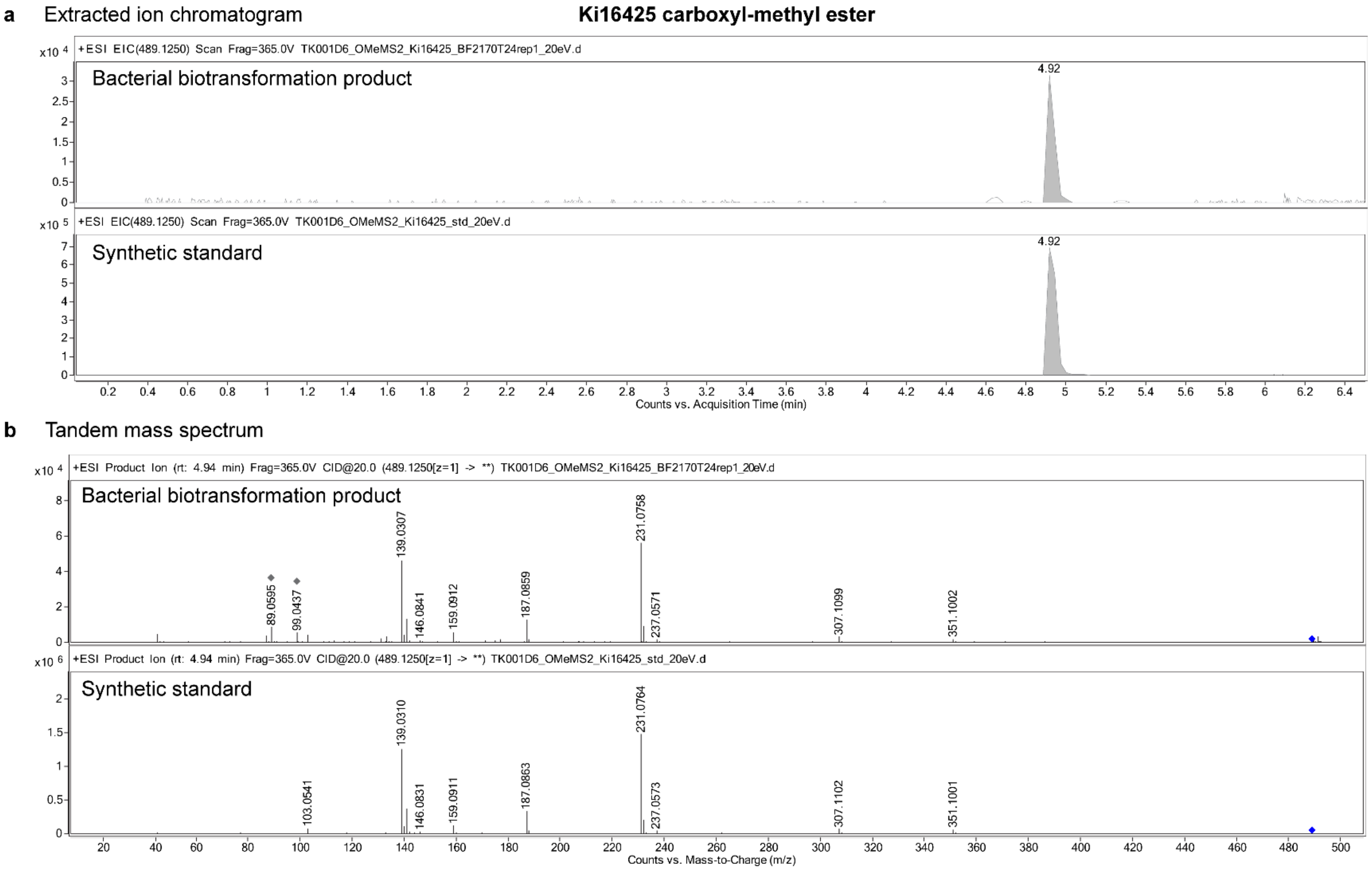

**Figure S7 | LC–MS/MS validation of Ki16425 carboxyl methyl ester produced by bacterial biotransformation.**

**a**, Extracted ion chromatograms and **b**, tandem mass spectra of the bacterial carboxyl-methylation product of Ki16425 and the synthetic standard. Blue diamond, precursor ion; grey diamond, background signals.

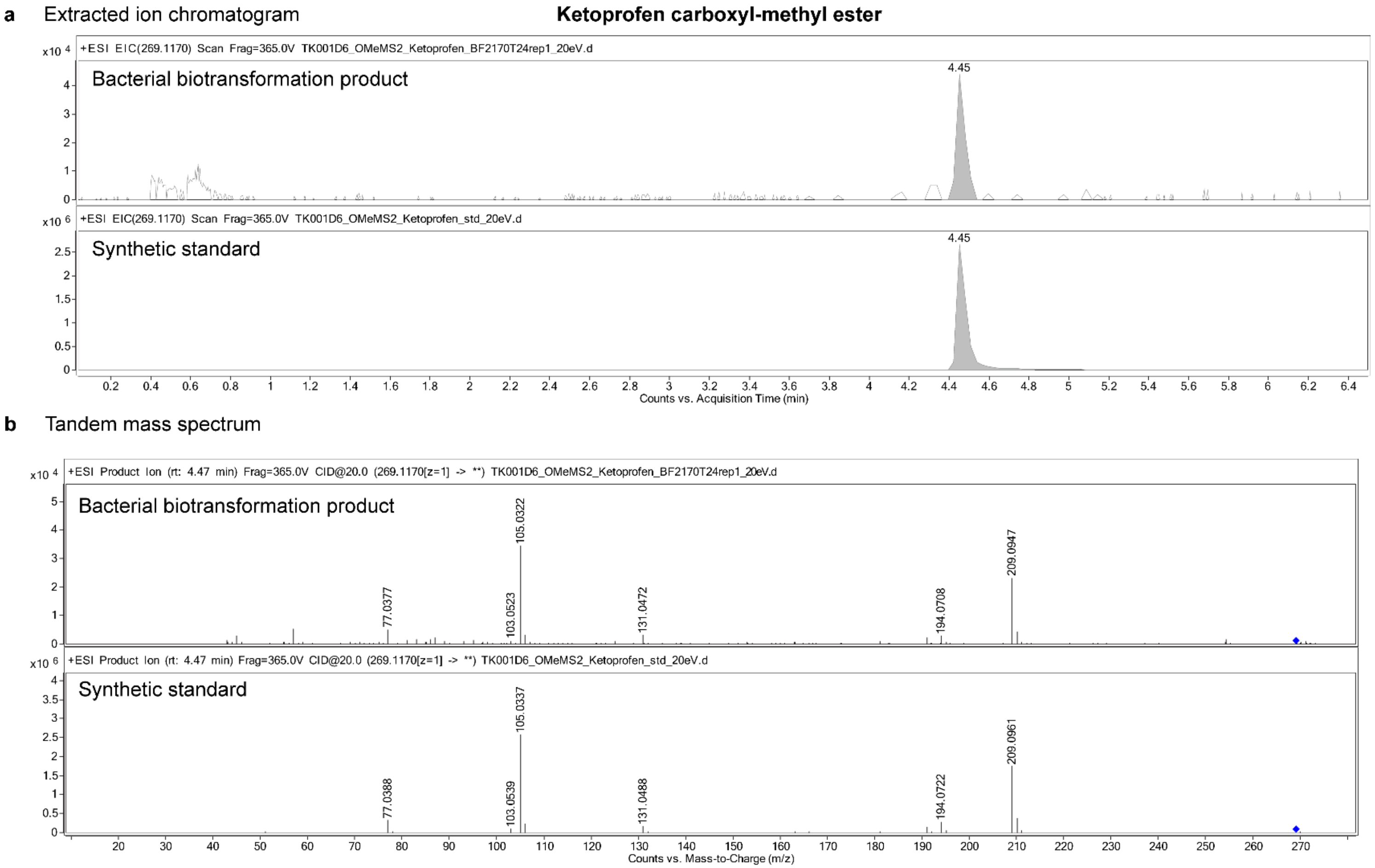

**Figure S8 | LC–MS/MS validation of ketoprofen carboxyl methyl ester produced by bacterial biotransformation.**

**a**, Extracted ion chromatograms and **b**, tandem mass spectra of the bacterial carboxyl-methylation product of ketoprofen and the synthetic standard. Blue diamond, precursor ion.

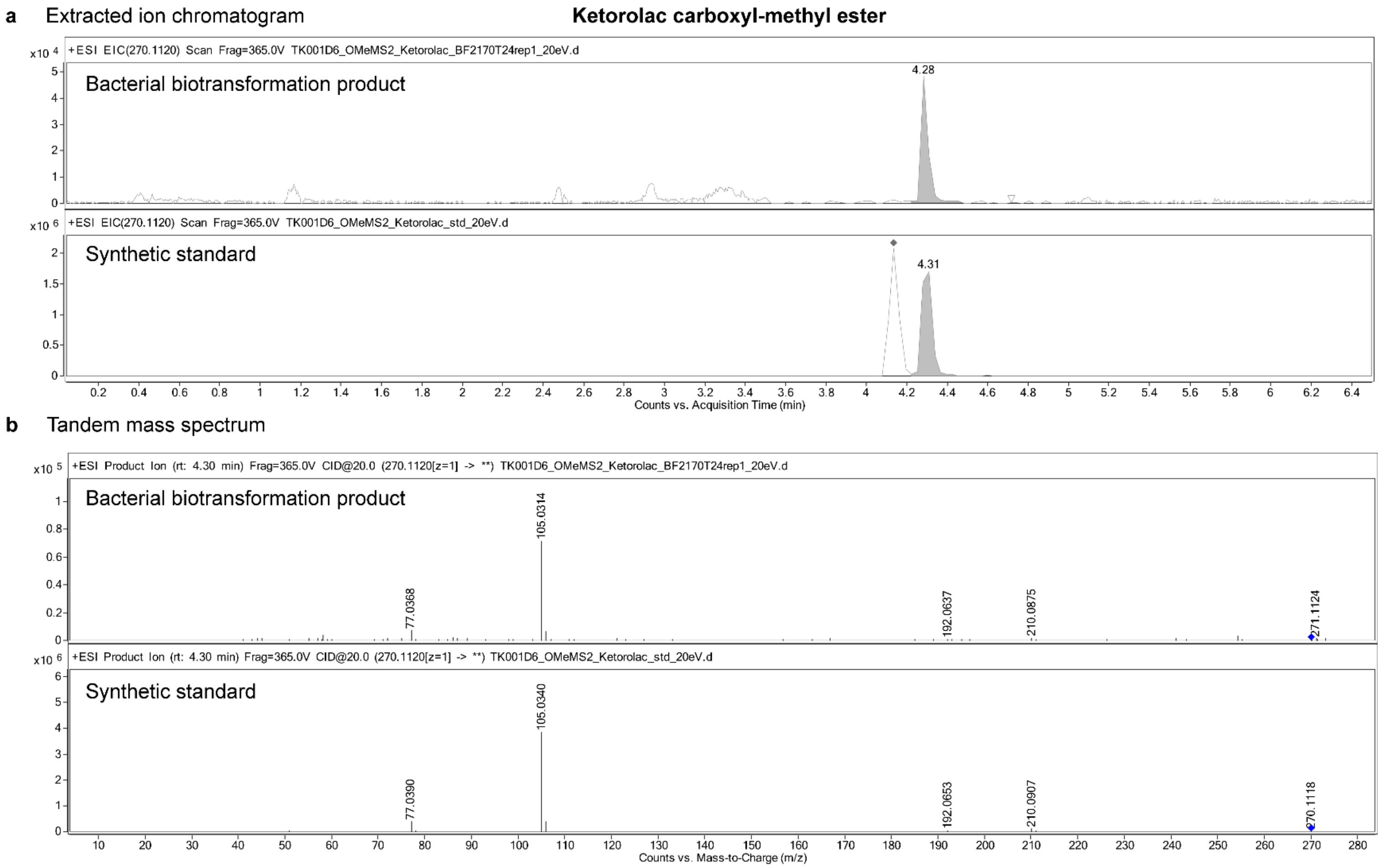

**Figure S9 | LC–MS/MS validation of ketorolac carboxyl methyl ester produced by bacterial biotransformation.**

**a**, Extracted ion chromatograms and **b**, tandem mass spectra of the bacterial carboxyl-methylation product of ketorolac and the synthetic standard. Blue diamond, precursor ion; grey diamond, background signals.

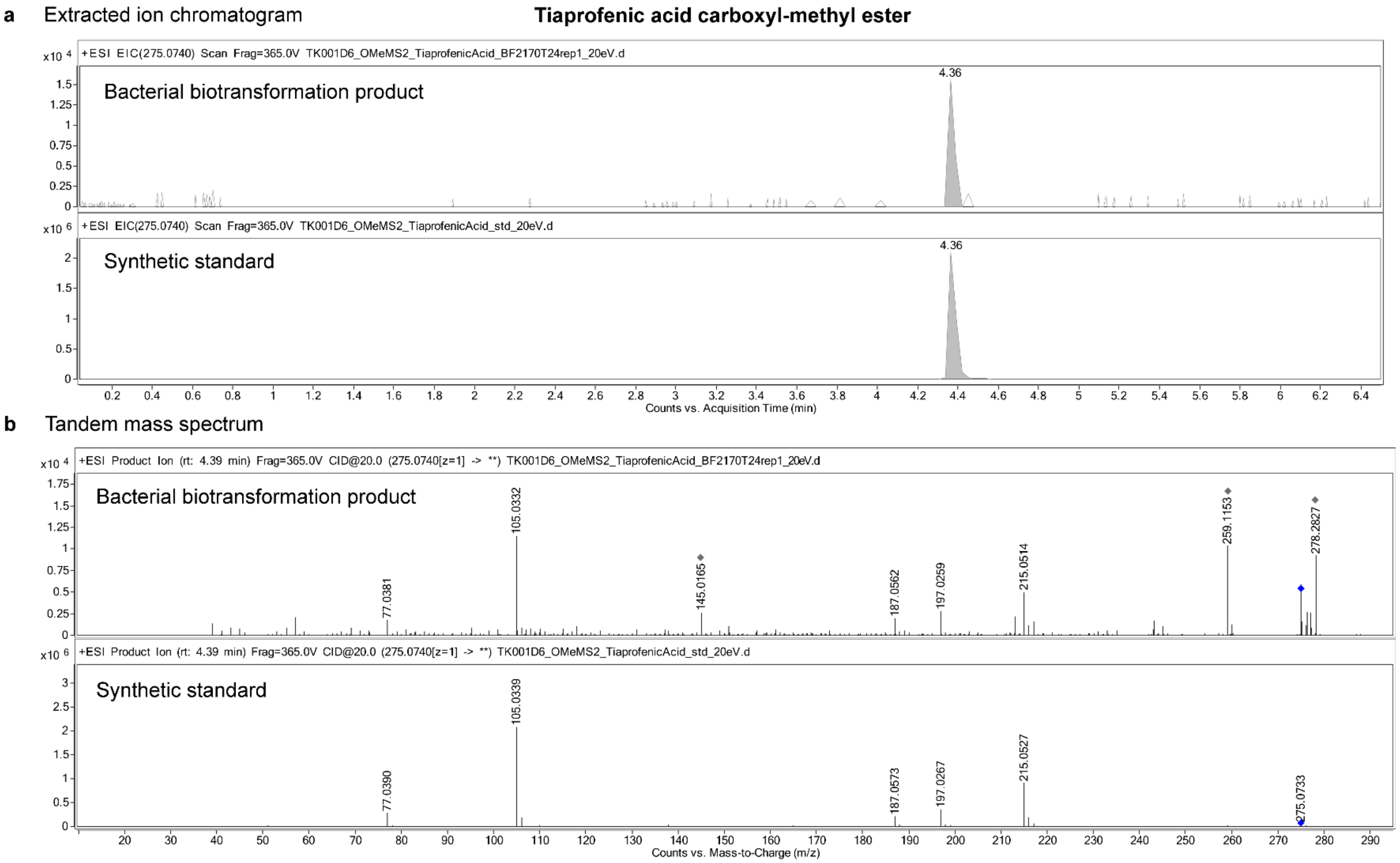

**Figure S10 | LC–MS/MS validation of tiaprofenic acid carboxyl methyl ester produced by bacterial biotransformation.**

**a**, Extracted ion chromatograms and **b**, tandem mass spectra of the bacterial carboxyl-methylation product of tiaprofenic acid and the synthetic standard. Blue diamond, precursor ion.

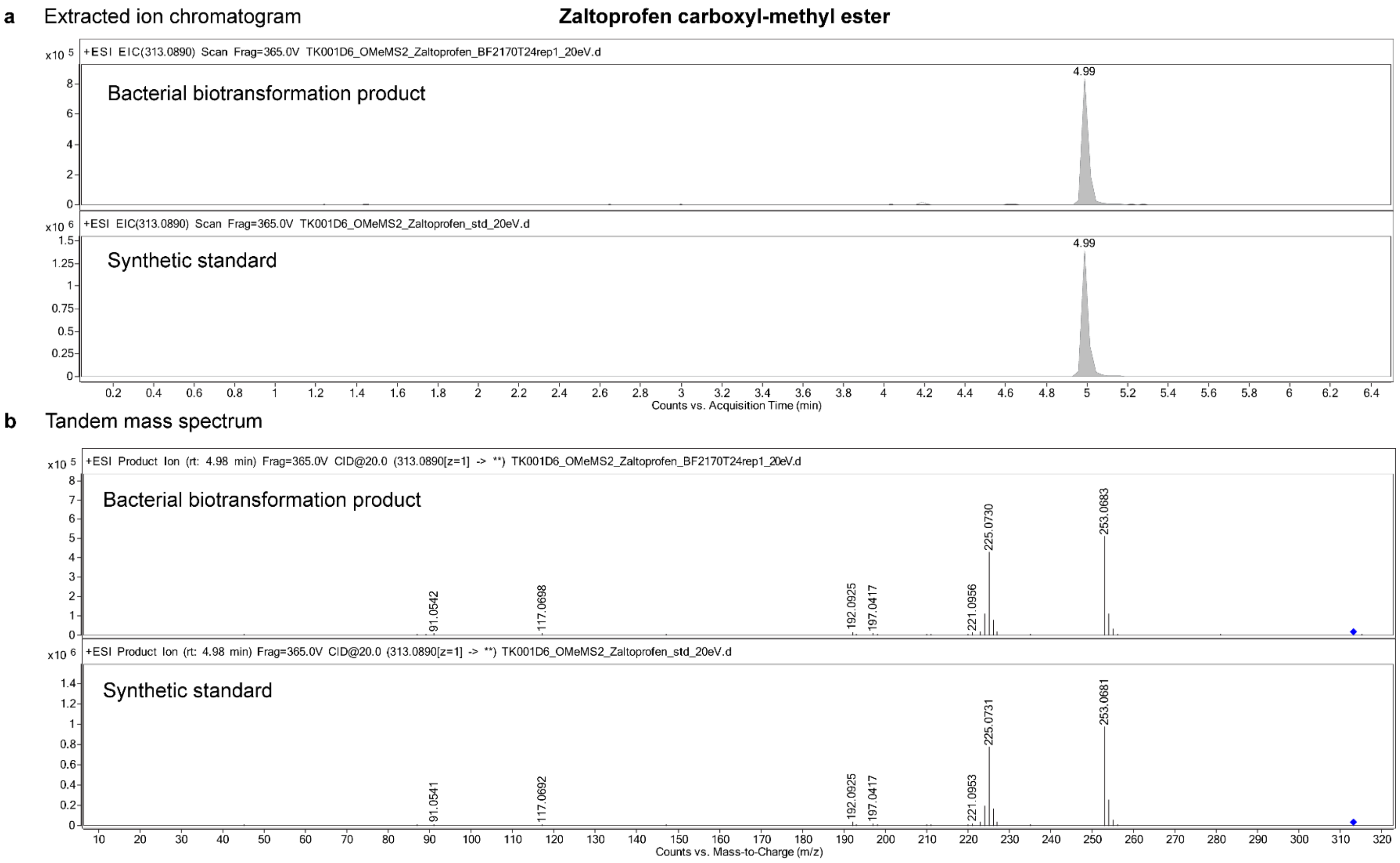

**Figure S11 | LC–MS/MS validation of zaltoprofen carboxyl methyl ester produced by bacterial biotransformation.**

**a**, Extracted ion chromatograms and **b**, tandem mass spectra of the bacterial carboxyl-methylation product of zaltoprofen and the synthetic standard. Blue diamond, precursor ion.

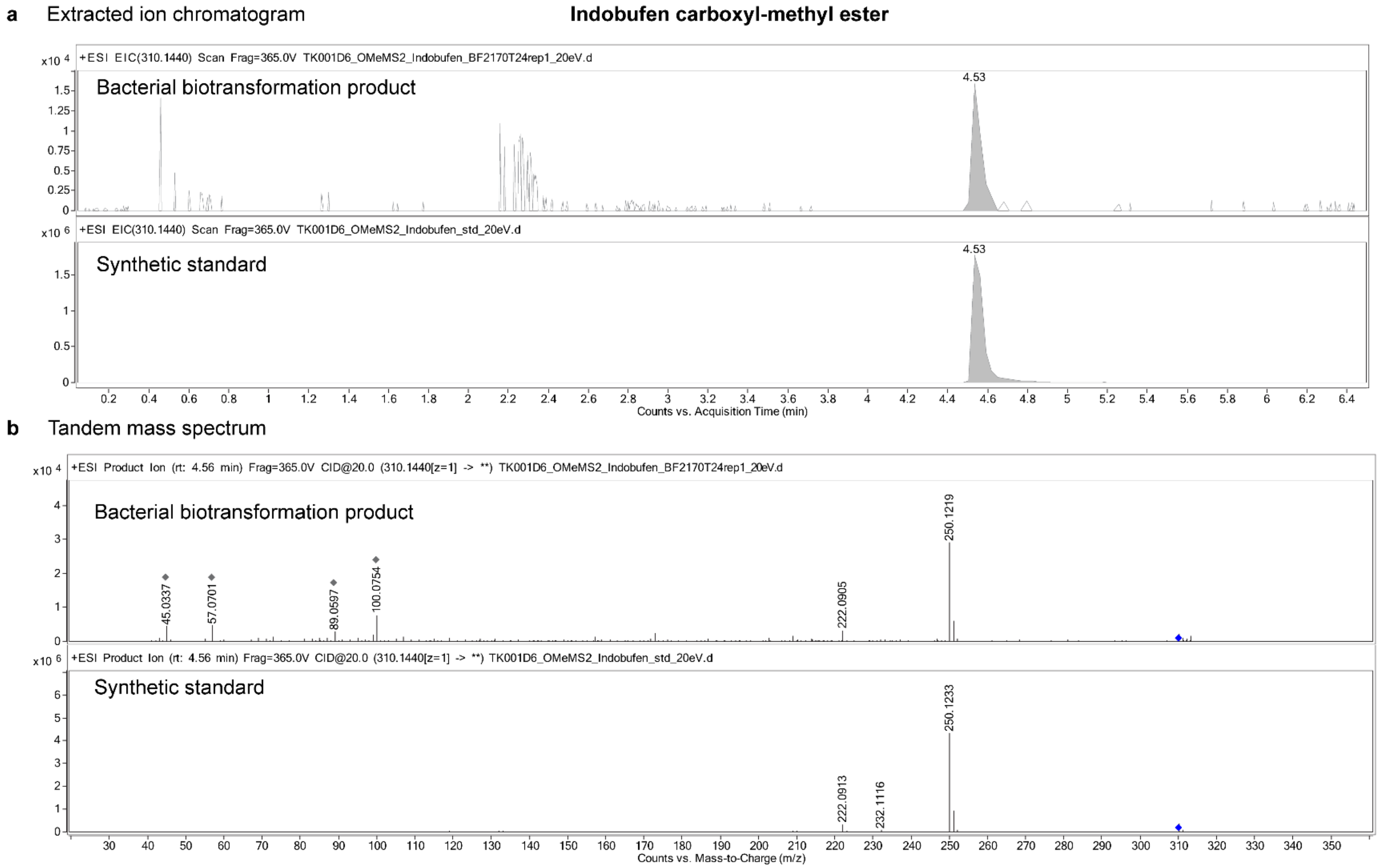

**Figure S12 | LC–MS/MS validation of indobufen carboxyl methyl ester produced by bacterial biotransformation.**

**a**, Extracted ion chromatograms and **b**, tandem mass spectra of the bacterial carboxyl-methylation product of indobufen and the synthetic standard. Blue diamond, precursor ion; grey diamond, background signals.

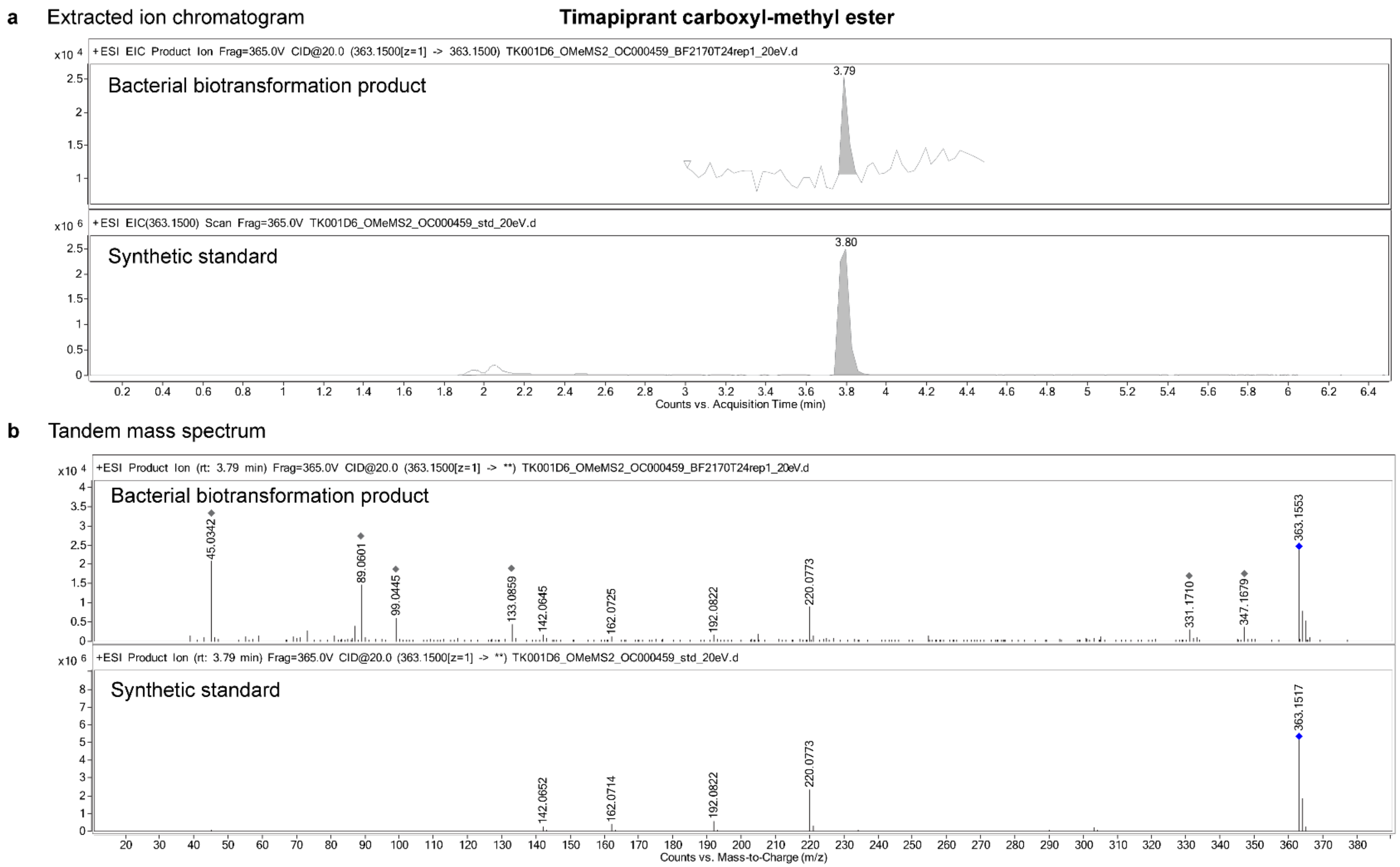

**Figure S13 | LC–MS/MS validation of timapiprant carboxyl methyl ester produced by bacterial biotransformation.**

**a**, Extracted ion chromatograms and **b**, tandem mass spectra of the bacterial carboxyl-methylation product of timapiprant and the synthetic standard. Blue diamond, precursor ion; grey diamond, background signals.

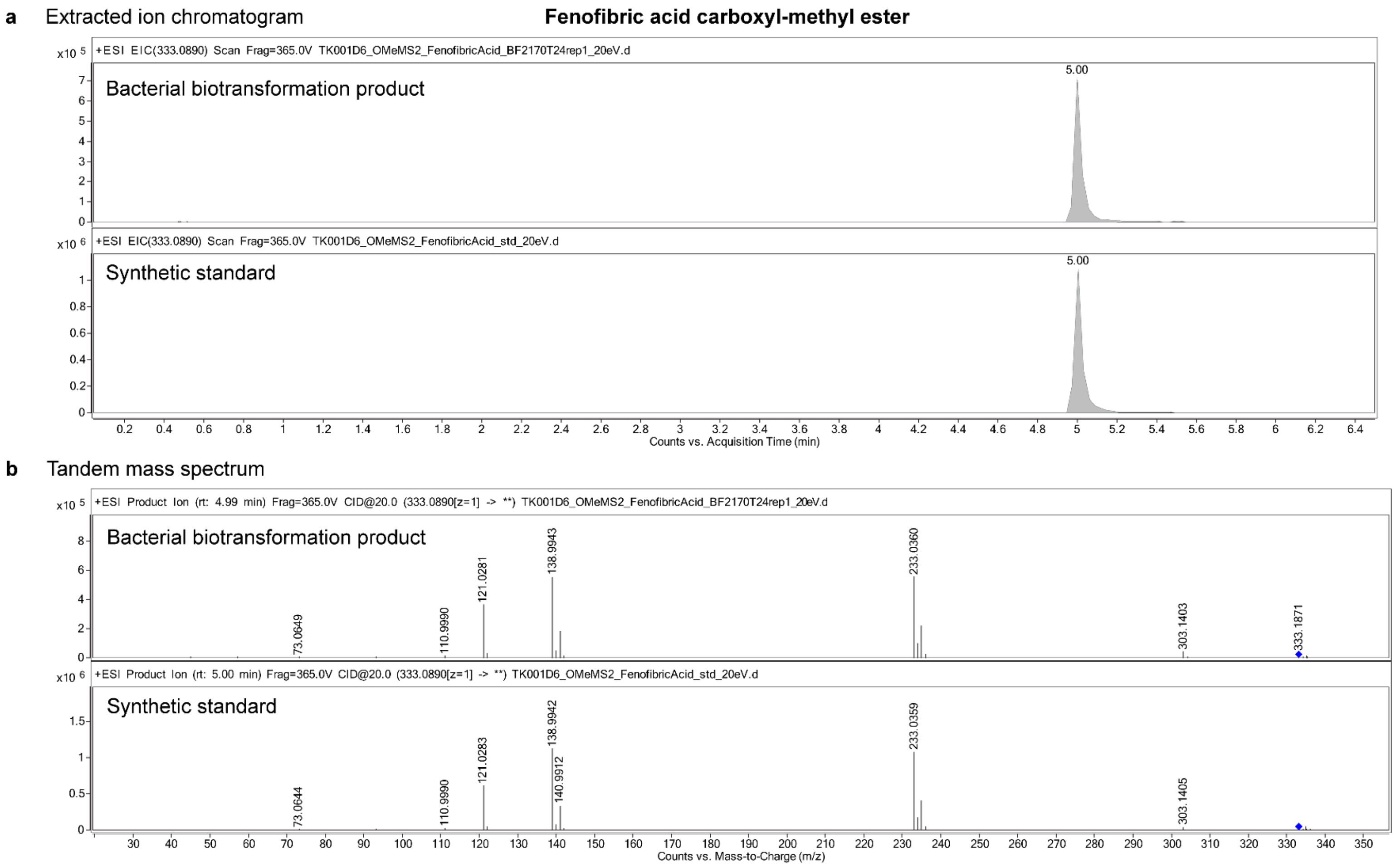

**Figure S14 | LC–MS/MS validation of fenofibric acid carboxyl methyl ester produced by bacterial biotransformation.**

**a**, Extracted ion chromatograms and **b**, tandem mass spectra of the bacterial carboxyl-methylation product of fenofibric acid and the synthetic standard. Blue diamond, precursor ion.
