## Supplementary Methods for "Gut bacteria generate prodrugs in situ increasing systemic drug exposure"

for

### Contents:

Additional methods for anaerobic cultivation, parameters for phylogenetic tree construction, protein structural prediction and alignment, chemical synthesis of carboxyl-methyl ester prodrugs, and detailed instrumental parameters used in high-throughput LC–QTOF and LC–QqQ analyses.

### Anaerobic cultivation

Anaerobic cultivation of bacteria was carried out in a vinyl anaerobic chamber (Coy Laboratory Products) equipped with a hydrogen sulfide removal column. The chamber was maintained with a gas mixture of 2% $H_2$, 12% $CO_2$, and 76% $N_2$, with the $O_2$ level maintained below 30 ppm. Culture media and agar plates were pre-reduced by incubating under anaerobic conditions for at least 16 h prior to use.

Growth curves of *Bacteroides fragilis* DSM2151 wild-type and Δ*bf2170* mutant strains were recorded under anaerobic conditions at 37 °C in 96-well conical-bottom plates (Thermo Scientific, 249952). Each well contained 100 µl of pre-reduced modified Gifu Anaerobic Medium (mGAM) inoculated with 1:10,000-diluted overnight cultures. Optical density at 578 nm was measured hourly using a microplate reader (BioTek Synergy H4).

### Parameters to construct phylogenetic trees

To reconstruct the UHGG v2.0.2 bacterial phylogeny, the bact120 protein alignment file was downloaded from the MGnify FTP server. IQ-TREE (v2.3.3) was used to generate a maximum-likelihood phylogenetic tree for the 4,716 bacterial species. The best-fit model was selected using the IQ-TREE ModelFinder with the -m MFP parameter. The LG+F+R10 model was selected based on the lowest Bayesian Information Criterion (BIC) score. Branch support was assessed using the Ultrafast Bootstrap approximation with 1,000 replicates (-B 1000). The phylogenetic tree was midpoint-rooted and visualized in iTOL (v7.2.1). A subtree representing the phylum *Bacteroidota* was extracted from the reconstructed UHGG bacterial phylogenetic tree using the R package ape.

### Protein structure prediction and alignment

Boltz-1 (v0.4.1) with ColabFold-based multiple sequence alignment with 10 recycling and 200 sampling steps and 5 diffusion samples was used to predict the structure of biomolecular complexes of bezafibrate ligand (CC(C)(C(=O)O)OC1=CC=C(C=C1)CCNC(=O)C2=CC=C(C=C2)Cl) and the selected methyltransferases. The diffusion samples with the highest confidence_score per complex were selected for further analysis. PyMOL (v3.1.5.1; Schrödinger, LLC) was used to visualize and superimpose the biomolecular complexes predicted with Boltz-1. RMSD values were calculated with "super" command.

### Chemical synthesis of carboxyl-methyl ester prodrugs

A single-step esterification protocol was employed to synthesize carboxyl methyl esters of carboxyl-containing drugs including BEZ, BEZ-$d_6$, efaproxiral, pranoprofen, zaltoprofen, orantinib, indobufen, timapiprant, and tiaprofenic acid. Synthesis of bromfenac carboxyl-methyl ester was not successful due to intra-molecular cyclization.

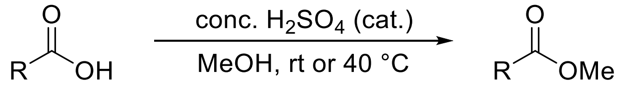


To a solution of a carboxylic acid drug (0.030 mmol) in methanol (2 ml) was added a catalytic amount of concentrated sulfuric acid (2 µl). The resulting mixture was stirred at room temperature overnight. In cases where the reaction proceeded slowly, the mixture was stirred at 40 °C. Upon completion, the solution was concentrated under reduced pressure. The crude residue was then dissolved in ethyl acetate and subsequently washed with saturated $NaHCO_3$ solution twice, followed by brine solution. The organic extract was dried over anhydrous $MgSO_4$ and filtered. All volatiles were removed under reduced pressure to afford the analytically pure methyl ester.

The purified products were structurally validated by $^1H$ and $^{13}C$ nuclear magnetic resonance spectroscopy on a Bruker Avance (400 MHz) NMR System at 298 K, and by UHPLC–MS analysis on an Agilent 1290 series equipment consisting of an Agilent 1290 quaternary pump, a 1290 sampler, a 1290 thermostated column compartment and a 1290 Diode array detector VL+ equipped with a quadrupole LC–MS 6120 and an Infinity 1260 ELSD. Compound purity was determined by ELSD monitoring.

**High-throughput LC–QTOF analysis: instrumental parameters in detail**

An Agilent Q–TOF 6546 mass spectrometer, or an Agilent 6550 Q–TOF mass spectrometer equipped with iFunnel technology, coupled to an Agilent 1290 Infinity II UHPLC dual-pump system was used to analyze samples from *in vitro* experiments.

A dual-column configuration was employed to provide high-throughput analysis, using two matched InfinityLab Poroshell 120 HPH-C18 columns (2.1 × 100 mm, 1.9 µm) maintained at 45 °C. A sample extract (5 µl) was injected with a sample flush-out factor of 10. Injection path cleaning was performed with 10 s of isopropanol (IPA) and 10 s of needle-washing (NW) solvent (water + 5% methanol + 0.1% formic acid). Chromatographic separation was performed by an analytical LC pump ('Binary Pump 2') using a linear gradient with solvent A (water + 0.1% formic acid) and solvent B (methanol + 0.1% formic acid), ramped from 5% to 95% B over 5.5 minutes and held at 95% B for 1 min at a flow rate of 0.6 ml/min. Meanwhile, the other column was regenerated offline using a secondary pump ('Binary Pump 1') with solvent A' (the same as NW solvent) and solvent B' (acetonitrile + 0.1% formic acid), applying a regeneration gradient held at 95% B' from 0 to 2 min, ramped to 0% B' by 3 min, and held at 0% B' until 6.5 min; a 0.5 min post-gradient re-equilibration was applied after column switching at 6.5 min, with the subsequent injection overlapped at 6.9 min (*i.e.*, 0.4 min post data acquisition). A 1/2-diluted IPA solution was used as the pump seal wash to prevent salt deposition and maintain lubrication.

A Dual AJS ESI source was operated with the following settings: gas temperature, 275 °C; drying gas, 13 l/min; nebulizer pressure, 40 psi; sheath gas, 12 l/min at 275 °C; Vcap, 3,500 V; and nozzle voltage, 2,000 V. Online mass calibration was performed using a reference solution containing purine ($[M+H]^+$ = *m/z* 121.0509) and hexakis(1H,1H,3H-perfluoropropoxy)phosphazene ($[M+H]^+$ = *m/z* 922.0098; CAS No. 58943-98-9), introduced via the secondary ESI source at a constant flow rate of 15 µl/min. Agilent MassHunter Workstation (v10.0) was used to acquire data. For LC–MS analysis, data were acquired in positive-ion mode over a mass range of *m/z* 100–1500 at 1.5 spectra/s in centroid format. For LC–MS/MS analysis, the targeted MS/MS mode with a preferred list for parent ions was used with an isolation width set to 'narrow (~1.3 *m/z*)', mass range of *m/z* 20-1500, delta RT of 2 min, and a fixed collision energy set to either 10, 20, or 40 eV in separate acquisitions with an acquisition speed of 3 spectra/s for MS and 1 spectrum/s for MS/MS. Agilent MassHunter Qualitative Analysis (v12.0) was used to examine chromatography and ion signals. Agilent MassHunter Quantitative Analysis (v10.0) was used for peak integration.

**LC–QqQ analysis: instrumental parameters in detail**

An Agilent 6495D Triple Quadrupole mass spectrometer equipped with iFunnel technology and coupled to an Agilent 1290 Infinity II UHPLC system (LC–QqQ) was used to quantify BEZ and its metabolites in samples from gnotobiotic mouse experiments. A single-column setup was used for all analyses. A sample extract (3 µl) was injected with a sample flush-out factor of 10. Injection path cleaning was performed with 10 s of acetonitrile and 10 s of methanol. Liquid chromatographic separation was performed using the same method as described above. To avoid carryover, blank injections were inserted between biological samples, and the overlapped injection was disabled. The ESI source was operated with the following parameters: gas temperature, 215 °C; drying gas, 15 l/min; nebulizer pressure, 25 psi; sheath gas, 12 l/min at 390 °C; capillary voltage, 3,500 V; and nozzle voltage, 0 V. Agilent MassHunter Workstation (v12.1) was used to acquire data in dynamic multiple reaction monitoring (dMRM) mode using quantifier and qualifier ion transitions as provided in Supplementary Table 25. Agilent MassHunter Qualitative Analysis (v12.0) was used to examine chromatography and ion signals. Agilent MassHunter Quantitative Analysis (v12.0) was used for peak integration.
